# TRIM52 is a primate-specific player in the DNA repair process under tight proteolytic control by a triad of giant E3 ligases

**DOI:** 10.1101/2024.05.16.594269

**Authors:** Alexandra Shulkina, Kathrin Hacker, Julian F. Ehrmann, Valentina Budroni, Ariane Mandlbauer, Johannes Bock, Daniel B. Grabarczyk, Luisa Cochella, Tim Clausen, Gijs A. Versteeg

## Abstract

Tripartite motif 52 (TRIM52) exhibits strong positive selection in humans, yet is lost in many other mammals. In contrast to what one would expect for such a non-conserved factor, *TRIM52* loss compromises cell fitness. We set out to determine the cellular function of TRIM52. Genetic and proteomic analyses revealed TRIM52’s involvement in resolving topoisomerase 2 (TOP2)-DNA cross-links, mitigating DNA damage and preventing cell-cycle arrest. Consistent with a fitness-promoting function, TRIM52 is upregulated in various cancers, prompting us to investigate its regulatory pathways. We found TRIM52 to be targeted for ultra-rapid proteasomal degradation by the giant E3 ubiquitin ligases BIRC6, HUWE1, and UBR4/KCMF1. BIRC6 mono-ubiquitinates TRIM52, with subsequent extension by UBR4/KCMF1. These findings underscore TRIM52’s pivotal role in DNA damage repair and regulation of its own abundance through multi-ligase degradation.

## Introduction

The evolutionary trajectory of organisms is often shaped by the interplay between genetic variation and environmental pressures. Understanding the function and regulation of positively-selected factors can provide insight into how different organisms adapt to their environment and thus how particular functions arose.

Evolutionary analyses have highlighted the significance of positive selection acting on genes associated with DNA damage repair, apoptosis regulation, and immune response pathways^1–4^. Particularly noteworthy among these are the tripartite motif (TRIM) protein E3 ligases, which have gained attention for their multifaceted roles in innate immunity, cellular homeostasis, and antiviral defense mechanisms^5^. Several members of the TRIM protein E3 ligase family have anti-retroviral functions and have co-evolved with the pathogens they counter^6^.

Interestingly, the *TRIM52* gene was acquired by the mammalian common ancestor, and evolved under positive selection pressure only in humans and some other primates^6^. In fact, it has been lost or pseudogenized in many other mammalian species. In contrast to what one would expect for such a non-conserved factor^6,7^, *TRIM52* ablation decreases cellular fitness and results in a p53-dependent decrease in proliferation^8,9^. Therefore, TRIM52 may play a role in resolution of DNA damage arising from cell-intrinsic DNA replication stress. In line with a cell fitness-promoting function, TRIM52 expression is upregulated in several cancers^10,11^.

This underpins the importance of understanding how intracellular abundance of TRIM52 is regulated, and what importance its regulation may have for its cellular function. In this context, we previously reported that TRIM52 is extremely rapidly turned-over by the proteasome, with a half-life of just 3-4 minutes^9^, positioning it as one of the most unstable proteins in the human proteome^12–16^.

TRIM52 has a unique, extended Really Interesting New Gene (RING) domain. The human genome encodes approximately 600 proteins with RING domains, a key feature of most ubiquitin E3 ligase enzymes^17^. It has remained unclear whether the unusual TRIM52 RING domain has E3 activity. However, mutagenesis experiments have shown that potential TRIM52 E3 ligase activity is dispensable for its own turn-over^9^, indicating that an unknown cellular machinery recognizes TRIM52 as a substrate, and marks it for degradation. This has positioned TRIM52 as an excellent model substrate to study the cellular and biochemical mechanisms by which cells mediate such rapid protein turnover.

Here we show that TRIM52 plays a role in resolution of topoisomerase 2 lesions stemming from cell-intrinsic processes such as transcription and genome replication. In the absence of *TRIM52*, covalent topoisomerase 2 retention on DNA lesion is increased, ultimately resulting in improper DNA break resolution, and downstream cell cycle arrest. Given that *TRIM52* is upregulated in several cancers^10,11^, we performed genetic and proteomic screens to identify its degradation machinery. This identified the giant E3 ligases BIRC6, HUWE1, and UBR4/KCMF1. Cell-based assays showed that these ligases mark TRIM52 for degradation through a highly acidic loop region in its RING domain. Mechanistically, *in vitro* experiments showed that TRIM52 degradation requires substrate recognition and ‘seeding’ of mono-ubiquitin by BIRC6, complemented by subsequent poly-ubiquitin chain extension by UBR4 and its co-factor KCMF1.

## Results

### TRIM52 functions in parallel to the non-homologous end joining DNA repair pathway

*TRIM52* ablation strongly diminishes cellular fitness in cell competition assays in various cell types^8,9,18^, mediated by aberrant p53 activation, and subsequent cell cycle arrest^8^. To gain insight into the cellular function of TRIM52 underlying this phenotype, we established a system to conduct an unbiased genetic screen for modifiers of *TRIM52* loss. We took advantage of a human RKO cell line (p53 wildtype colon carcinoma) harboring an inducible Cas9 cassette that has been effectively used in various genome-wide CRISPR screens (Fig. 1a)^19,20^. When we introduced sgRNAs targeting *TRIM52* and induced Cas9 expression, we observed practically complete depletion of TRIM52 protein and markedly diminished cellular fitness in a competitive proliferation assay (Fig. 1a-c).

**Figure 1-.**
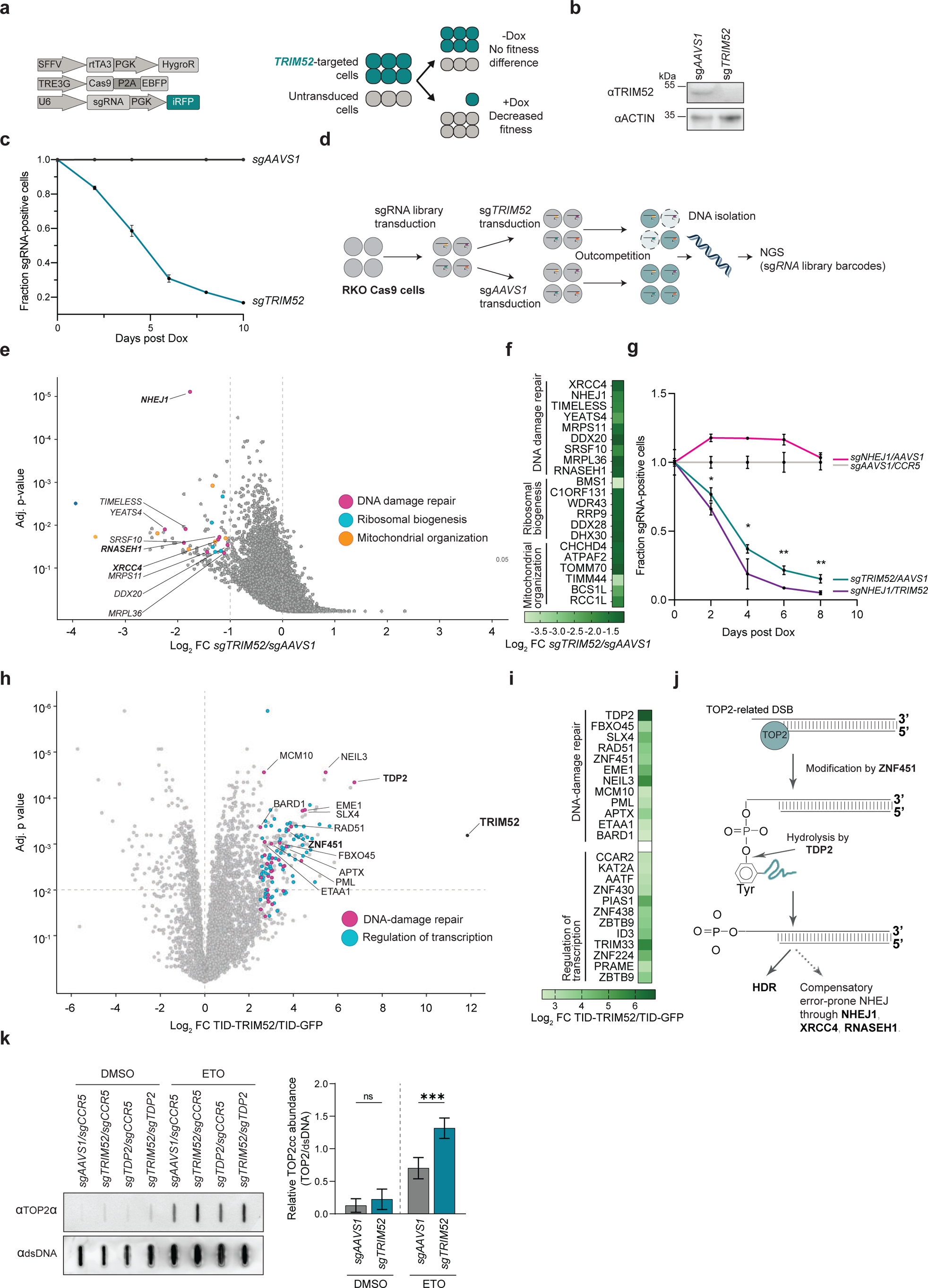
TRIM52 is required for resolution of TOP2 DNA damage lesions. (**a**) Schematic representation of the lentiviral expression vectors used to modify RKO Cas9 cells for knock-out and cell competition assays. (**b**) RKO cells harboring Dox-inducible Cas9 were transduced with sgRNA vectors targeting *TRIM52* or the safe-harbor locus *AAVS1*. Cas9 was induced by Dox treatment for 5 days. Whole cell lysates (WCL) were analyzed by WB. (**c**) Relative cell fitness of sg*TRIM52*-transduced cells compared to untransduced cells was determined by measuring the percentage of iRFP fluorescent cells over the indicated period of time and normalized to the fraction of sg*AAVS1*-transduced cells relative to untransduced cells. (Data represent biological samples, n = 3). (**d**) Schematic representation of genetic modifier screen. (**e**) RKO cells expressing Cas9 and transduced with a lentiviral genome-wide sgRNA library were further transduced with individual sgRNAs targeting *TRIM52* or *AAVS1*. Following Cas9 induction, cells were grown for 12 doublings and integrated sgRNA-coding sequences were determined by NGS analysis of gDNA. Genes isolated from the population are plotted as Log2 fold change relative to the sgRNA library representation determined in the unselected cell population. Factors on the left-hand side of the plot are identified as genes which lead to synthetic lethality upon ablation in *TRIM52*-targeted cells. Non-*TRIM52*-specific genes were filtered out by comparison with data from cells with *DGCR8* KO. Factors involved in DNA-damage repair are marked in pink and labelled by name. (**f**) Filtered *TRIM52* ablation-specific genes were selected and their Log2 fold change compared to *AAVS1* plotted in a heatmap and grouped by their function. (**g**) Relative cell fitness of sg*TRIM52*, sg*NHEJ1* and sg*TRIM52*/sg*NHEJ1* transduced cells compared to untransduced cells was determined by measuring the percentage of iRFP fluorescent cells over the indicated period. sg*TRIM52* fractions were normalized to sg*AAVS1/CCR5*-transduced cells, relative to untransduced cells. (Data represent biological samples, n = 3). sg*TRIM52/AAVS1* and sg*NHEJ1*/TRIM52 samples were analyzed by multiple unpaired t-tests. ns: p > 0.05, *: p < 0.05, **: p < 0.01, ***: p < 0.001) (**h**) A TurboID-TRIM52 fusion was expressed, cells treated with proteasome inhibitor for 5 h., and biotin for the last 15 min. Biotinylated proteins were purified under denaturing conditions, and analyzed by mass-spectrometry. Factors involved in DNA-damage repair are marked in pink and labelled by name. Data represent biological samples; n = 3. (**i**) Putative interactors of TRIM52 with p-value < 0.05 and Log2 fold change > 2.5 were selected, analyzed by gene ontology enrichment analysis, and top hits per ontology category plotted in a heatmap by their Log2 fold change enrichment compared to the TurboID-EGFP control. (**j**) Schematic representation of TOP2 cleavage complex resolution. (**k**) RKO-Cas9 cells transduced with sgRNAs targeting safe harbor loci *AAVS1*/*CCR5, TRIM52*, *TDP2* or both were treated with Dox for 5 days to induce Cas9. Samples were harvested, and subjected to RADAR analysis. Isolated protein-DNA covalent complexes were analyzed by slot blot, quantified and normalized to dsDNA levels. (Data represent biological samples, n = 4. Data were analyzed by 2-way ANOVA. ns: p > 0.05, *: p < 0.05, **: p < 0.01, ***: p < 0.001).

We performed the genome-wide modifier screen in RKO cells harboring a Dox-inducible Cas9-P2A-EBFP expression cassette and a genome-wide lentiviral sgRNA library, targeting one gene per cell. Into this complex cell population, we transduced a lentivirus expressing two validated sgRNAs targeting *TRIM52*, or two sgRNAs targeting the safe harbor locus *AAVS1*, as a control (Fig. 1d, Extended data Fig.1a). This enabled identification of synthetically lethal genes whose mutation enhanced the *TRIM52*-loss-associated fitness defects. To identify *TRIM52*-specific modifiers, we performed a parallel counter-screen in cells with a knockout of *DGCR8*, a key component of the microRNA processing machinery, as its ablation reduces cell fitness to a similar extent as ablation of *TRIM52*, but it is expected to act via unrelated mechanisms.

The screen did not identify any specific factors that increased cell fitness upon ablation (Supplementary data 1). However, it did identify genes whose ablation was specifically synthetically lethal with *TRIM52* loss, but not with *DGCR8* knock-out (Fig. 1e-f). Several high confidence genetic interactors were associated with the error-prone, non-homologous end-joining (NHEJ) DNA damage response (Fig. 1e-f, Supplementary data 1; *NHEJ1*, *XRCC4*, and *RNASEH1*), indicating that this pathway can in part compensate for functional *TRIM52* loss. In line with these findings, depletion of *NHEJ1* (a central factor in this DNA repair pathway) by itself did not decrease cell fitness in a competition assay, while targeting *NHEJ1* in the context of *TRIM52* loss diminished cell fitness more than depletion of *TRIM52* alone (Fig. 1g). From these results we concluded that the cellular role of TRIM52 can in part be functionally compensated by the error-prone NHEJ1-dependent DNA repair pathway, and thus that TRIM52 likely acts in a parallel pathway to NHEJ. This begins to explain our previous observation that loss of *TRIM52* elicits p53 activation, which can be triggered by cell-intrinsic DNA damage^21–23^.

### TRIM52 is required for resolution of TOP2-mediated DNA lesions

To gain further insight into which aspect of DNA damage and repair TRIM52 is involved in, we set out to identify physical interactors of TRIM52 using TurboID proximity labeling^24^ (Extended Data Fig. 1b). We generated RKO cells with a cassette for Dox-inducible expression of a TurboID-TRIM52 fusion, or TurboID-EGFP as a control^24^. After careful titration of Dox to achieve comparable protein levels (Extended data Fig. 1c), we treated these cells with biotin for 15 min to label proteins in proximity. We performed labeling in the presence of proteasome inhibitor to stabilize the pool of likely DNA-associated TRIM52, which is otherwise low due to rapid proteasomal turn-over (Extended data Fig. 1d). Under these conditions, TRIM52 localized to nuclear puncta (Extended data Fig. 1e), reminiscent of previously reported DNA repair complexes^25^. We then isolated biotinylated proteins under denaturing conditions and used nano liquid chromatography coupled mass-spectrometry (nLC-MS/MS) for identification.

We observed a strong enrichment in proteins related to the DNA damage response and transcription regulation (Fig. 1h-i, Extended data Fig. 1f, Supplementary data 2). Transcription can lead to DNA damage through various mechanisms, such as the formation of R-loops, and collisions between transcription and DNA replication machineries^26,27^. Pathway enrichment analysis of the proteins involved in the DNA damage response revealed a strong enrichment for factors involved in homology-dependent repair (HDR; Extended data Fig. 1g-h (inner circle), and Supplementary data 3, adj. p < 0.0001). This suggested that TRIM52 either plays a functional role in **i)** HDR in general, or **ii)** a specific DNA damage sensing pathway that ultimately recruits and activates the HDR machinery.

Two of the strongest identified TRIM52 interactors (Fig. 1h-i, and Extended data Fig. 1g,h), Tyrosyl-DNA Phosphodiesterase 2 (TDP2) and Zinc Finger Protein 451 (ZNF451), are not part of the core HDR machinery, but rather play key roles upstream of HDR in specifically sensing and resolving topoisomerase 2 (TOP2) lesions^28–32^. TOP2 resolves topological problems in DNA that originate from DNA replication, transcription, and chromosome dynamics^33^. TOP2 introduces DSBs to reduce torsional stress, and becomes covalently linked to the 5’ phosphate via a tyrosine residue^28–30^. For error-free repair of these DSBs, TOP2 must be removed from DNA, which is achieved through the action of TDP2 and the SUMO E3 ligase ZNF451 that facilitates TDP2 hydrolase activity on stalled TOP2 cleavage complexes^28–32^. This leaves 5’ phosphates free for repair by HDR and NHEJ (Fig. 1j)^34–36^. These activities are critical for cell fitness, as their inhibition results in p53 activation, and cell cycle arrest even in the absence of exogenous DNA damage^28,29,31,37^, as seen upon loss of *TRIM52*^8^. Therefore, we hypothesized that TRIM52 plays a role in the repair of TOP2-associated DSBs, playing an important function in the TDP2-dependent HDR pathway (Fig. 1j).

To test this hypothesis, we measured TOP2-DNA covalent adducts using the ‘Rapid Approach to DNA Adduct Recovery’ (RADAR) assay^38,39^. In this assay, cellular DNA with covalently associated protein complexes is precipitated, after which TOP2 levels are measured by slot blot. We reasoned that if TRIM52 plays a functionally important role in resolution of TOP2 lesions, its ablation would sensitize cells to TOP2 toxins, such as Etoposide (ETO). We exposed cells in which either the safe-harbor loci *AAVS1/CCR5* or *TRIM52* were targeted with sgRNAs to sub-saturating concentrations of ETO, then measured TOP2 cleavage complexes (TOP2cc) associated with DNA by RADAR assay. Under untreated conditions, *TRIM52*-targeting showed a trend for increased DNA-associated TOP2cc complexes (Fig. 1k). ETO treatment increased the levels of TOP2cc by 5.5-fold in sg*AAVS1/CCR5* control cells, which was further increased ∼2-fold in *TRIM52*-targeted cells (Fig. 1k), indicating the requirement of TRIM52 for efficient hydrolysis of TOP2-DNA phospho-tyrosyl bonds.

Taken together, these results show that TRIM52 plays a role in the resolution of TOP2-dependent DNA lesions. These findings help explain previous observations showing that loss of *TRIM52* results in cell-intrinsic activation of p53, and ultimately cell cycle arrest^8^.

### TRIM52 is degraded by the ubiquitin-proteasome system

Considering TRIM52’s function in DNA repair, and the fact that its levels are upregulated in several cancers^10,11^, we next set out to understand how its abundance is regulated. First, to determine which cellular degradation pathways contribute to TRIM52 turn-over, we treated RKO cells expressing Ollas-tagged EGFP-TRIM52 with the proteasome inhibitor epoxomicin, the lysosome inhibitor bafilomycin A, or the neddylation inhibitor MLN-4924, which inhibits Cullin ubiquitin E3 ligases. We then analyzed Ollas-EGFP-TRIM52 protein levels in whole cell lysates (WCL) by Western blot (WB) and flow cytometry. Proteasome inhibition by epoxomicin increased Ollas-EGFP-TRIM52 protein levels by 2.5-fold in WB (Fig. 2a, and Extended data Fig. 2a), and 3.6-5.2-fold by flow cytometry (Extended data Fig. 2b). None of the other inhibitors affected TRIM52 concentrations, indicating that it is predominantly degraded through proteasomal degradation.

**Figure 2-.**
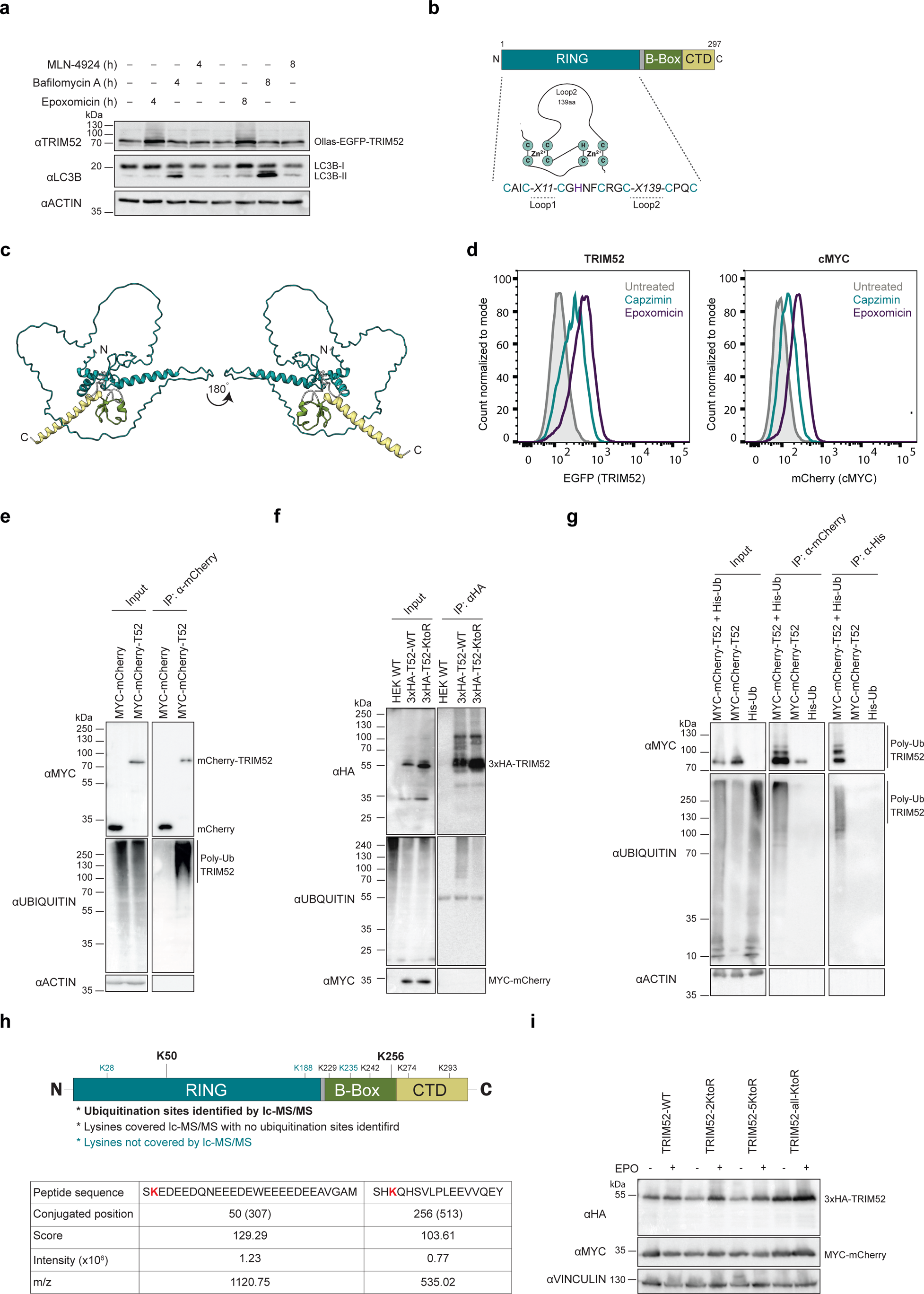
TRIM52 is degraded by the ubiquitin-proteasome system. (**a**) RKO cells expressing Ollas-tagged EGFP-TRIM52 were treated for 4 or 8 h. with the proteasome inhibitor epoxomicin, the lysosome inhibitor bafilomycin A, or the neddylation inhibitor MLN-4924. WCLs were analyzed by WB. (**b**) Schematic representation of TRIM52 domain organization. (**c**) AlphaFold2 model for TRIM52 structure prediction. (**d**) RKO cells expressing Ollas-tagged EGFP-TRIM52 or mCherry-cMYC fusion proteins were treated for 5 h. with the proteasomal 20S catalytic core particle inhibitor epoxomicin, or 19S regulatory particle-specific inhibitor capzimin. Ollas-EGFP-TRIM52 and mCherry-cMYC protein levels were determined by measuring EGFP or mCherry fluorescence using flow cytometry, and their mean fluorescence intensity (MFI) plotted. (**e**) RKO cells expressing MYC-tagged mCherry, or MYC-tagged mCherry-TRIM52 were treated with epoxomicin for 5 h., lysed under denaturing conditions, RFP-trap IPs performed, and analyzed by WB for endogenous ubiquitin. (**f**) HEK-293T cells transfected with 3xHA-tagged TRIM52 or TRIM52 in which all lysines were mutated to arginines were treated with epoxomicin for 5 h., lysed under denaturing conditions, TRIM52 was immunoprecipitated, and its ubiquitination analyzed by WB. (**g**) HEK-293T cells expressing MYC-tagged mCherry or mCherry-TRIM52, and His-tagged ubiquitin were treated with epoxomicin for 5 h., dual-affinity purified by RFP-trap and NiNTA. Eluates were analyzed by WB, and analyzed by nLC-MS/MS for the identification of ubiquitination sites. (**h**) Identified ubiquitination sites plotted on schematic TRIM52 domain representation. (**i**) HEK-293T cells were transfected with 3xHA-tagged TRIM52 WT and TRIM52 lysine-to-arginine mutants, treated with epoxomicin for 5 h., and lysates analyzed by WB.

TRIM52 has a long, low complexity loop 2 within its RING domain (Fig. 2b) that is predicted to be unstructured by AlphaFold2.0 (Fig. 2c, Extended data Fig. 2c-d). To test whether TRIM52 is degraded by 20S or 26S proteasomes, we compared the effect of the 19S regulatory particle inhibitor capzimin (which exclusively inhibits 26S proteasomes)^40^ with the 20S core particle inhibitor epoxomicin (which inhibits both 20S and 26S proteasomes). Since capzimin is a less effective inhibitor than epoxomicin^40^, we compared its effects on TRIM52 to cMYC, which is degraded in a ubiquitin- and 26S proteasome-dependent manner^41,42^. We treated cells expressing Ollas-tagged EGFP-TRIM52 or mCherry-cMYC fusion proteins with the indicated proteasome inhibitors, then analyzed the effects on their steady-state levels by flow cytometry. Both inhibitors comparably increased TRIM52 and cMYC levels (Fig. 2d and Extended data Fig. 2e). Epoxomicin increased TRIM52 levels by 1.7-fold and cMYC by 2.7-fold. As expected, the effect of capzimin was less pronounced (Fig. 2d, and Extended data Fig. 2e): 0.5-fold for TRIM52 and 0.75-fold for cMYC. Since the relative effects of both inhibitors were comparable between both substrates, we concluded that, like cMYC, TRIM52 is predominantly degraded in a 26S proteasome-dependent manner.

To test whether TRIM52 is ubiquitinated in cells, we immunoprecipitated MYC-tagged mCherry-TRIM52 from RKO cell lysates and checked for associated poly-ubiquitin chains by WB. Consistent with 26S proteasomal degradation (Fig. 2d, and Extended data Fig. 2e), MYC-mCherry-TRIM52 was strongly ubiquitinated, whereas a MYC-tagged mCherry control was not detectably modified (Fig. 2e). Lysine is the most prominent residue for post-translational modification by ubiquitin. Therefore, we compared ubiquitination of HA-tagged WT TRIM52 with a mutant in which all lysines were mutated to arginines. Consistent with reduced degradation, steady-state levels of the lysine-less TRIM52 mutant were increased relative to its WT counter-part, whereas its ubiquitination was reduced by 70% (Fig. 2f, Extended data Fig. 2f).

To identify the sites of ubiquitination in TRIM52, we tandem-purified MYC-tagged mCherry-TRIM52 from cells expressing his-tagged ubiquitin (Fig. 2g) and performed nLC-MS/MS identification of peptides with di-Gly ubiquitin remnants^43,44^. This revealed a strong presence of di-Gly residues on ubiquitin itself on K11, K29, K33, K48, and K63, with K48 being >10 fold as abundant as any of the other detected linkages (Supplementary data 4), indicating K48-linked chains to be the major poly-ubiquitin modification on TRIM52. In addition, we identified two high confidence di-Gly sites on TRIM52: one site within its loop 2 region of the RING domain (K50) and another in the BBox domain (K256) (Fig. 2h; Supplementary data 4). We detected peptides covering four of the remaining lysines (K229, K242, K276, K293) by nLC-MS/MS, but they were not significantly ubiquitinated. No peptides containing K28, K188, and K235 were identified by nLC-MS/MS, for which it thus remained unclear whether they were ubiquitinated (Fig. 2h) and could contribute to degradation.

To test the functional importance of these lysine residues for TRIM52 degradation, we tested sensitivity to proteasome inhibition for TRIM52 mutants in which groups of lysines were mutated to arginines. Mutation of the two identified lysines (2KtoR; K50/K256) did not result in TRIM52 stabilization, nor did additional mutation of the three lysines that were not covered by nLC-MS/MS analysis (5KtoR; Fig. 2i). Only mutation of all lysines (all-KtoR) increased steady-state TRIM52 levels, and rendered it insensitive to epoxomicin treatment (Fig. 2i).

Together, these results show that TRIM52 is K48 poly-ubiquitinated at a minimum of two predominant lysine residues. Removal of these sites was not sufficient to prevent TRIM52 degradation, suggesting that the E3 ligase machinery targeting TRIM52 can modify non-dominant sites in their absence.

### TRIM52 is targeted for degradation by multiple giant E3 ligases

To identify the factors responsible for ultra-rapid degradation of TRIM52, we performed another genetic screen in the RKO-Cas9-EBFP cell line. In this case, we introduced a stably expressed dual protein stability reporter (Fig. 3a). This bicistronic reporter was designed to translate stable MYC-tagged mCherry and unstable Ollas-tagged EGFP-TRIM52 in equimolar amounts through a P2A ribosomal skip site^45^, yet accumulate low EGFP-TRIM52 steady-state levels as a result of its degradation (Fig. 3a).

**Figure 3-.**
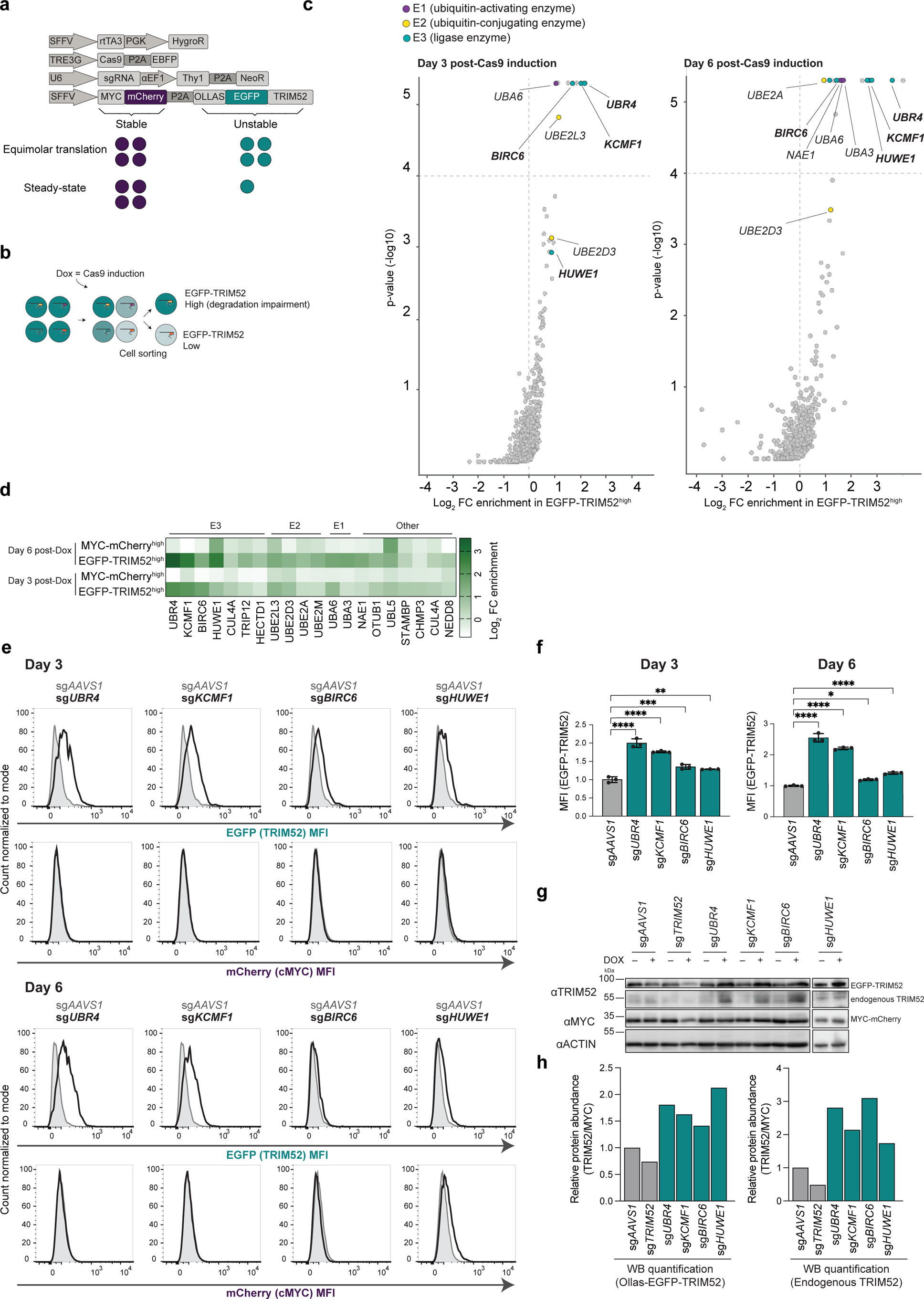
TRIM52 is targeted for degradation by multiple giant E3 ligases. (**a**) Schematic representation of the expression vectors in the screening cell line. mCherry and EGFP-TRIM52 are expressed in an equimolar manner, yet EGFP-TRIM52 accumulates at low steady-state levels, resulting from its rapid proteasomal turn-over. (**b**) Schematic representation of the genetic screen to identify TRIM52 regulators. EGFP^high^ cells with potential knock-outs in factors involved in TRIM52 degradation were collected by FACS, and their integrated sgRNA CDSs quantified by NGS, relative to those from unsorted cell pools. (**c**) Screening cells were transduced with an sgRNA library targeting ubiquitin-related genes and treated with Dox for 3 or 6 days. Cells expressing the highest and lowest 1-2% of each fluorophore were collected by FACS, their integrated sgRNA CDSs quantified by NGS, and differential abundance plotted. (**d**) Heatmap displaying the Log2 fold change for genes enriched in the EGFP-TRIM52^high^ sorted population after exclusion of genes enriched in mCherry^high^ population on days 3 and 6. (**e**) RKO-Cas9 cells expressing MYC-mCherry-P2A-Ollas-EGFP-TRIM52 (teal) or mCherry-cMYC-P2A-EBFP (purple) as a control were transduced with lentiviral vectors expressing the indicated sgRNAs. Cas9 expression was induced for 6 days, after which EGFP-TRIM52 and mCherry-cMYC protein levels were quantified by flow cytometry, and **(f)** MFI plotted (Data represent biological samples, n = 3. Data were analyzed by 1-way ANOVA. ns: p > 0.05, *: p < 0.05, **: p < 0.01, ***: p < 0.001, ****: p < 0.0001)., or **(g)** analyzed by WB, and **(h)** quantified by densitometry.

We selected a monoclonal cell line that had sufficiently high EGFP-TRIM52 levels to enable screening by flow cytometry, and yet mirrored endogenous TRIM52 instability. Specifically, we determined that levels of EGFP-TRIM52, but not mCherry, were increased by proteasome inhibition (Extended data Fig. 3a). In addition, the turn-over of EGFP-TRIM52 measured upon translation inhibition was dramatically faster than that of mCherry (Extended data Fig. 3b-c). Even though overexpression did extend the half-life of the reporter to 30 min, compared to 3.3 min for endogenous TRIM52, this is still a very unstable protein. For reference, endogenous cMYC is considered highly unstable with a half-life of 15-30 min^12^. Therefore, we concluded it to be a suitable reporter for screening. Given that we established a new monoclonal line, we also confirmed the efficiency and inducible control of Cas9-mediated gene editing in this cell line (Extended data Fig. 3d).

To screen for factors that induce the degradation of TRIM52, we transduced a lentiviral sgRNA-encoding library into our reporter cell line at a low multiplicity of infection to ensure targeting of only one gene per cell (Fig. 3b, Extended data Fig. 3e). The sgRNA library was designed to target ∼1000 genes, including E1 activating enzymes, E2 conjugases, E3 ligases, deubiquitinating enzymes, and ubiquitin-interacting proteins. At two time points after Cas9 induction (3 and 6 days), we used FACS to select the top 1-2% EGFP^high^ cells, with potential defects in TRIM52 degradation. In parallel, we collected a comparable control cell pool for mCherry to exclude genes that generally increase protein abundance (Fig. 3b and Extended data Fig. 3f). The sgRNA loci present in these various cell populations were amplified, quantified by NGS, and plotted relative to their representation in unsorted cell pools (Fig. 3c-d).

This screen resulted in strong enrichment of a select group of E1, E2 and E3 ligases in the EGFP-TRIM52^high^ cells that were absent in the mCherry^high^ control population (Fig. 3c-d, Extended data Fig. 3g). Specifically, we identified two E1 activating enzymes, UBA3 and UBA6, the E2 conjugating enzymes UBE2D3, UBE2L3, UBE2A and UBE2M, and three high-confidence giant E3 ligases: BIRC6, HUWE1 and UBR4 with its known interactor KCMF1^46^ (Fig. 3c-d, Supplementary data 5). We validated the effect of the three identified E3 ligases in independently generated polyclonal cell pools (Fig. 3e-f). Moreover, to test their relative specificity for controlling TRIM52 turn-over, we targeted the same E3 ligases in cells expressing an mCherry-tagged version of the proteasomal-targeted transcription factor cMYC. In contrast to the effect on EGFP-TRIM52, knock-out of *UBR4/KCMF1* or *BIRC6* had no measurable influence on the steady-state levels of mCherry-cMYC (Fig. 3e; rows 2 and 4). This was consistent with similar genetic screens that identified degradation-regulators of cMYC or another unrelated transcription factor, IRF1^47^. HUWE1 has been previously identified as a regulator of cMYC degradation^47,48^, and consistent with these findings, its knock-out did moderately increase cMYC protein levels, especially at day 6 post Cas9 induction (Fig. 3e). However, we still included HUWE1 as a candidate TRIM52 regulator considering its broad substrate range^49,50^.

To confirm that these candidate E3 ligases are *bona fide* regulators of endogenous TRIM52, and not only of the TRIM52 reporter used for genetic screening, we also showed that loss of the E3 ligases increased endogenous TRIM52 levels (Fig. 3g-h). To test whether the increased TRIM52 levels indeed stemmed from increased protein stability, we targeted the individual E3 ligases in RKO cells stably expressing HA-tagged TRIM52 and an internal MYC-tagged mCherry control, and determined protein levels by WB at different timepoints after protein synthesis inhibition by CHX (Extended data Fig. 4a). The half-life of HA-TRIM52 in the sg*AAVS1* control was 11.8 min and increased 3-5 fold to 31-60 min in the different E3 ligase knock-outs (Extended data Fig. 4b). Taken together, we identified BIRC6, HUWE1, and UBR4/KCMF1 as E3 ligases that functionally mediate TRIM52 protein turn-over.

**Figure 4-.**
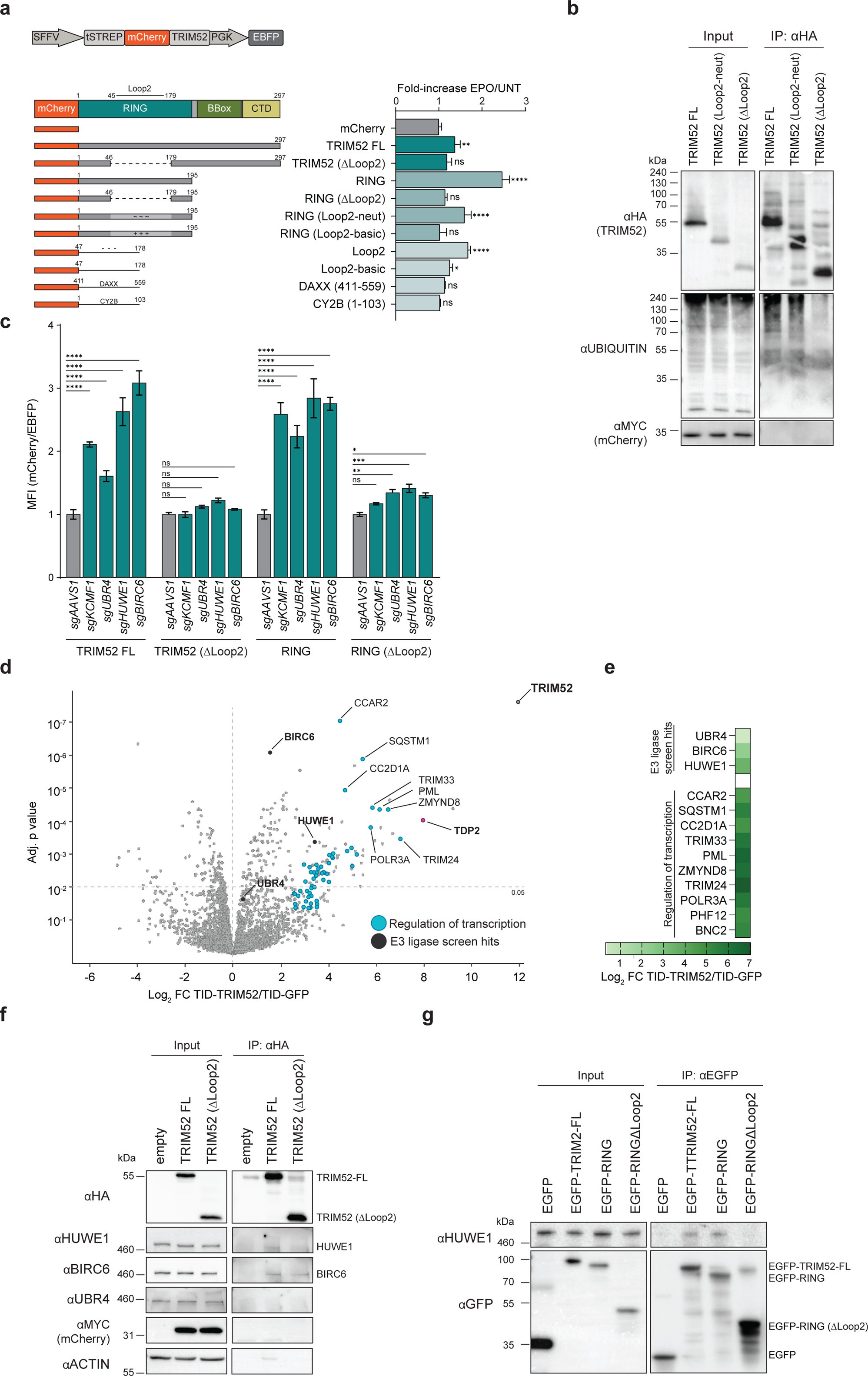
BIRC6, HUWE1 and UBR4/KCMF1 target the extended loop 2 region in the TRIM52 RING domain. (**a**) HEK-293T cells were transfected with plasmids expressing the indicated TwinStrep-mCherry-tagged TRIM52 expression constructs and an internal EBFP as a control. Cells were treated with proteasomal inhibitor epoxomicin for 5 h., after which protein levels were quantified by flow cytometry, and normalized to EBFP levels and relative to corresponding untreated samples and plotted (Data represent biological samples, n = 3. Data were analyzed by 1-way ANOVA. ns: p > 0.05, *: p < 0.05, **: p < 0.01, ***: p < 0.001, ****: p < 0.0001). (**b**) HEK-293T cells were transfected to express HA-tagged WT TRIM52, TRIM52 neutral loop2 (Loop2-neut) or TRIM52 in which the loop 2 region is reduced to the size of other RING proteins (ΔLoop2), treated with epoxomicin for 5 h., TRIM52 immunoprecipitated, and ubiquitination levels analyzed by WB. (**c**) HEK-293T cells constitutively expressing Cas9 and sgRNAs targeting the indicated genes were transfected with plasmids expressing TwinStrep-mCherry-tagged TRIM52, TRIM52 (ΔLoop2), RING, or RING (ΔLoop2), as well as an internal control EBFP. Protein levels were quantified by flow cytometry, normalized to EBFP and the sg*AAVS1* control and plotted (Data represent biological samples, n = 3. Data were analyzed by 2-way ANOVA. ns: p > 0.05, *: p < 0.05, **: p < 0.01, ***: p < 0.001, ****: p < 0.0001). (**d**) RKO cells expressing a TurboID-TRIM52 fusion protein were incubated with biotin for 15 min., after which biotinylated proteins were purified under denaturing conditions, and analyzed by mass-spectrometry. Data represent biological samples; n = 3. (**e**) Heatmap displaying the Log2 fold change of selected interactors of TRIM52. (**f**) HEK-293T cells expressing 3xHA-tagged TRIM52 were treated with epoxomicin for 5 h., after which TRIM52 was immunoprecipitated, and analyzed by WB for co-immunoprecipitation with HUWE1, BIRC6, and UBR4. (**g**) Purified TwinStrep-tagged EGFP-TRIM52, EGFP-RING and EGFP-RINGΔLoop2 were incubated with purified human HUWE1 for 3 h. Twin-Strep-EGFP-TRIM52 was then immunoprecipitated form the samples using EGFP-trap beads and analyzed for complex formation with HUWE1 by WB.

### BIRC6, HUWE1, and UBR4/KCMF1 target the extended loop 2 region in the TRIM52 RING domain

Having identified E3 ligases that mediate TRIM52 turn-over, we next set out to identify which TRIM52 protein domains and features confer its instability, as they may provide insight into how the identified E3 ligases recognize TRIM52 as a substrate. To this end, we transfected HEK-293T cells with vectors expressing mCherry-tagged WT TRIM52, TRIM52 mutants, or individual protein domains in isolation (Fig. 4a). We gated cells for similar levels of the independently expressed EBFP internal control (Fig. 4a). As a measure of the instability of the tested proteins, we treated these cell pools with proteasome inhibitor then determined the levels of their mCherry-tagged fusion proteins by flow cytometry. Consistent with our previous findings that TRIM52 is rapidly degraded, mCherry-TRIM52 significantly increased upon epoxomicin treatment (Fig. 4a) compared to the mCherry control. Instability was lost upon removal of the extended loop 2 region in the TRIM52 RING domain (ΔLoop2), indicating this region to be the main determinant of TRIM52 degradation. Consistent with this notion, the RING domain in isolation was sufficient to confer instability to mCherry, but not in the absence of loop 2 (Fig. 4a; compare samples 4 and 5 to mCherry).

To test whether the highly acidic composition of the loop 2 region is required for TRIM52 turn-over, we mutated acidic D/E residues to neutral or basic residues in a manner that is predicted to maintain the disordered nature of the loop 2 region. Conversion of the loop 2 region into a more basic and positively charged variant completely stabilized the fusion protein (Fig. 4a). Likewise, a RING mutant carrying a neutral loop 2 was more stable than the WT RING domain as evidenced by a significant reduction in its stabilization upon proteasome inhibition (Fig. 4a). We then investigated whether the acidic/negative unstructured loop 2 region of TRIM52 was sufficient to confer instability in isolation. Indeed, fusion of just loop 2 to mCherry rendered it sensitive to proteasome inhibition, whereas a basic variant, an unstructured region from CY2B, or a region of similar length and acidic amino acid composition from DAXX did not confer instability (Fig. 4a and Extended data Fig. 5a). We conclude that the loop 2 region in the TRIM52 RING is the major determinant of TRIM52 degradation, and likely a stand-alone degron. While its biased amino acid composition is required for its destabilizing effect, additional features beyond charge and disorder likely play a role, as a comparable region from DAXX did not confer instability (Fig. 4a and Extended data Fig. 5a).

**Figure 5-.**
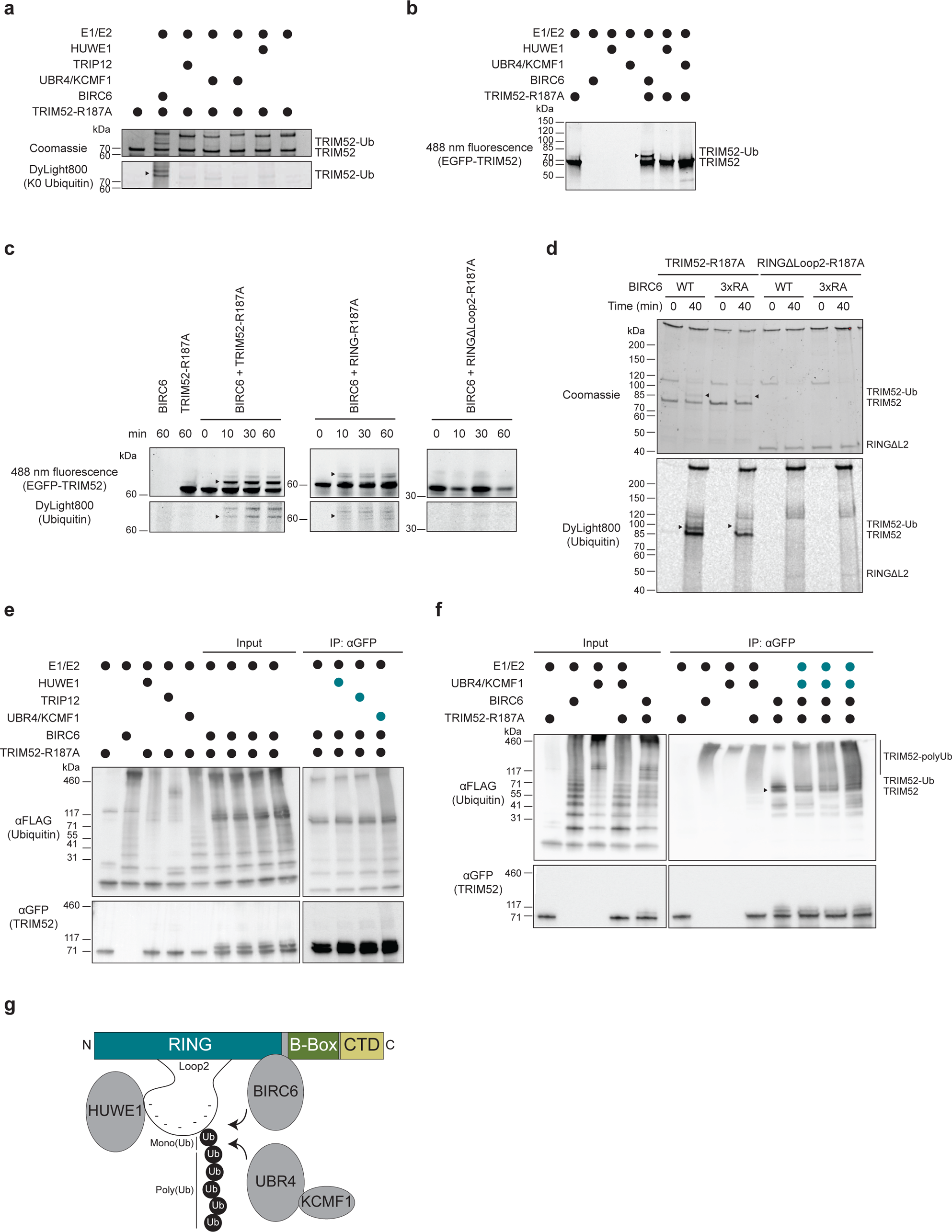
UBR4/KCMF1 extend poly-ubiquitin chains on sites directly mono-ubiquitinated by BIRC6. (**a**) *In vitro* ubiquitination assay of purified EGFP-TRIM52-R187A with recombinant E3 ligases, their cognate E1 and E2 enzymes and DyLight800-labeled K0 ubiquitin. The reactions were incubated for 1h. at 37 °C in the presence of ATP. (**b**) *In vitro* ubiquitination assay of EGFP-TRIM52-R187A with BIRC6 and in combination with HUWE1 or UBR4/KCMF1, their cognate E1 and E2s and WT ubiquitin. The reactions were incubated for 1h. at 37 °C in the presence of ATP. (**c**) Time-course ubiquitination assay of EGFP-TRIM52-R187A, EGFP-RING-R187A, and EGFP-RING (ΔLoop2) with recombinant BIRC6, UBA6, and DyLight800-labeled ubiquitin. The reactions were incubated for indicated times at 37 °C in the presence of ATP. (**d**) Ubiquitination assay of EGFP-TRIM52-R187A, EGFP-RING (ΔLoop2) with BIRC6-WT or BIRC6-3xRA, UBA6 and DyLight800-labeled ubiquitin. The reactions were incubated for indicated times at 37 °C in the presence of ATP. (**e**) Ubiquitination assay coupled to immunoprecipitation. Left: ubiquitination assays of EGFP-TRIM52-R187A with BIRC6, HUWE1, TRIP12, and UBR4/KCMF1, their cognate E1 and E2s as well as FLAG-labeled ubiquitin; right: αGFP IP of the indicated samples: the immunoprecipitated EGFP-TRIM52-R187A was further incubated with the indicated E3 ligases (labeled in teal), their cognate E1 and E2s as well as FLAG-ubiquitin. The reactions were incubated for 1 h. at 37 °C in the presence of ATP. (**f**) Ubiquitination assay coupled to immunoprecipitation. Input: ubiquitination assays of EGFP-TRIM52-R187A with BIRC6 or UBR4/KCMF1, their cognate E1 and E2s and FLAG-labeled ubiquitin. IP: αGFP IP of the indicated samples: the immunoprecipitated EGFP-TRIM52-R187A was further incubated in ubiquitination reactions with UBR4/KCMF1, their cognate E1 and E2s (labeled in teal) as well as FLAG-ubiquitin. The reactions were incubated for 1 h. at 37 °C in the presence of ATP. (**g**) Schematic model of TRIM52 degradation.

To test whether TRIM52 ubiquitination is dependent on its loop 2 region, we isolated HA-tagged TRIM52 from cells and analyzed its ubiquitination by WB. Indeed, full-length HA-TRIM52 was strongly ubiquitinated, while almost all ubiquitination was lost in the absence of loop 2 (Fig. 4b). In agreement with an intermediate phenotype, a TRIM52 variant with a neutral loop 2 was still partially ubiquitinated (Fig. 4b), consistent with the finding that it was still partially unstable (Fig. 4a, sample 6). To test whether the RING loop 2 is also the region through which the E3 ligases mediate TRIM52 degradation, we ablated the individual E3 ligases in HEK-293T cells expressing either full-length WT TRIM52, a TRIM52-ΔLoop2 mutant, only the RING domain, or the RING domain lacking the loop 2 region. Ablation of the identified E3 ligases (Extended data Fig. 5b) significantly increased the levels of full-length TRIM52 and its RING domain in isolation by 1.6-3.1 fold (Fig. 4c, set 1 and 3, and Extended data Fig. 5c), while neither full-length TRIM52 nor the RING domain in isolation were stabilized upon removal of the loop 2 region (Fig. 4c, set 2 and 4). Therefore, both TRIM52 ubiquitination and degradation of TRIM52 by the identified giant E3 ligases are dependent on the loop 2 region.

### BIRC6 and HUWE1 bind to TRIM52

We reasoned that there are three main possibilities for how the loop 2 region could contribute to TRIM52 turn-over: **i)** it is the site of E3 binding/recognition, **ii)** it is the site of ubiquitination, or **iii)** it is the site for both E3 binding and ubiquitination. To distinguish between the first two options and the last option, we utilized the finding that fusion of loop 2 to mCherry rendered this otherwise stable fluorophore in part unstable (Fig. 4a). We thus predicted that if a given E3 ligase either binds or ubiquitinates outside of loop 2, an mCherry-loop2 construct would not be affected upon E3 ligase ablation (either option 1 or 2). However, if a given E3 mediates TRIM52 targeting in a loop 2-autonomous manner (option 3), then such a construct should be stabilized in the absence of the E3 ligase.

To test this, we ablated the E3 ligases in cells expressing either mCherry or mCherry-loop2 and determined the protein levels by flow cytometry. As expected, none of the knock-outs influenced mCherry levels (Extended data Fig. 5d). However, *UBR4* and *HUWE1* ablation significantly increased mCherry-loop2 levels (Extended data Fig. 5d), indicating that loop 2 alone is sufficient for targeting by these two E3s. In contrast, loss of *BIRC6* did not significantly affect mCherry-loop2 (Extended data Fig. 5d), suggesting that BIRC6 either binds TRIM52 in loop 2 but ubiquitinates it outside of this region, or *vice versa*.

Genetic interactors identified in our screen (Fig. 3c-d) could affect TRIM52 turn-over **i)** directly by binding it as a substrate, **ii)** acting as ubiquitin chain extending E4 enzymes on (mono-) ubiquitinated substrates, or **iii)** indirectly by affecting other proteins. We reasoned that cellular complex formation between TRIM52 and any of the identified E3 ligases would indicate that TRIM52 is recognized as a direct substrate. Lack of such cellular complex formation would be consistent with possible an E4 function or indirect effects, yet would not rule out TRIM52 as a direct substrate as some interactions are transient.

To identify the E3 ligases interacting with TRIM52 in an unbiased manner, we performed additional TurboID proximity labeling experiments^24^. In contrast to the TurboID samples generated for assessing the functional role of TRIM52 (Fig. 1h), these samples were not generated in the presence of proteasome inhibitor, in order to maximize capture of the total steady-state interactome. Consistent with the earlier TurboID data (Fig. 1h-j, and Supplementary data 2), BIRC6 and HUWE1 were significantly enriched in TurboID-TRIM52 samples, compared to TurboID-EGFP controls (Fig. 4d-e, and Extended data Fig. 5e). While UBR4 peptides were detected in TurboID-TRIM52 samples, they were not significantly enriched over the TurboID-EGFP control (Fig. 4d-e). KCMF1 peptides were not detected in the TurboID-TRIM52 samples at all. Consistent with a role of TRIM52 in resolution of DNA damage lesions stemming from cell intrinsic processes (Fig. 1h-i), we also identified interactors involved in DNA-dependent transcription (Fig. 4d-e, Extended data Fig. 5f).

Together, these findings indicate that BIRC6 and HUWE1 likely form a complex with TRIM52 in cells, making them strong candidates for E3 ligases directly ubiquitinating TRIM52. Although the absence of detectable UBR4/KCMF1 interaction does not exclude their direct ubiquitination of TRIM52, it suggested that a complex of UBR4/KCMF1 could contribute to TRIM52 degradation in an E4 ligase capacity through ubiquitin chain extension, or another indirect manner.

Results presented in Extended data Fig. 5d indicated that HUWE1 likely binds inside loop 2 and ubiquitinates inside of it, whereas BIRC6 either binds inside or outside loop 2. To test this by co-IP, we isolated full-length TRIM52 or its Δloop2 mutant from cells and analyzed the association of HUWE1 and BIRC6. Full-length TRIM52 did in fact co-IP both HUWE1 and BIRC6 (Fig. 4f-g). While HUWE1 binding was strongly reduced in TRIM52-ΔLoop2 samples, BIRC6 association was not substantially changed. These results are consistent with a model in which HUWE1 binds and ubiquitinates inside loop 2, whereas BIRC6 likely binds the TRIM52 RING outside of loop 2, yet ubiquitinates within it.

### UBR4/KCMF1 extend poly-ubiquitin chains on sites directly mono-ubiquitinated by BIRC6

To further investigate how the identified E3 ligases operate together to ubiquitinate TRIM52, we used *in vitro* ubiquitination assays. TwinStrep-EGFP-TRIM52 (TRIM52 hereafter) was expressed in insect cells and isolated by streptavidin affinity purification (Extended Data Fig. 6a). Recombinant TRIM52 protein auto-ubiquitinated and synthesized free poly-ubiquitin chains in *in vitro* reactions (Extended Data Fig. 6b), from which we concluded that TRIM52 itself has E3 ligase activity, despite its extended RING loop 2 region. Since auto-ubiquitination is dispensable for TRIM52 turn-over in cells^9^, and the goal was to investigate TRIM52 as a substrate of the identified E3 ligases, we aimed to exclude ubiquitination by TRIM52 itself. We therefore expressed and purified a catalytically inactive TRIM52 allosteric linchpin^51^ mutant (TRIM52-R187A), which no longer synthesized poly-ubiquitin chains (Extended Data Fig. 6c).

To test which E3 ligases could ubiquitinate TRIM52, and how many individual residues are modified within the substrate, we incubated TRIM52-R187A with the identified E3 ligases, ATP, and their cognate E1 and E2 enzymes. To assess the number of modified residues, we included lysine-less Dylight800-labeled ubiquitin (K0 ubiquitin), which prevents background signal from poly-ubiquitin chain formation. BIRC6 ubiquitinated TRIM52 on three residues in a time-dependent manner (Fig. 5a, and Extended Data Fig. 6d-e), whereas UBR4/KCMF1 and HUWE1 did not efficiently ubiquitinate TRIM52-R187A (Fig. 5a). Similar reactions with wild-type ubiquitin yielded similar results (Fig. 5b), indicating that BIRC6 multi-mono-ubiquitinates full-length TRIM52. There was no evidence for efficient *in vitro* ubiquitination by UBR4/KCMF1 or HUWE1.

Cell-based experiments had indicated that the TRIM52 RING domain is sufficient as a substrate, and the loop 2 region is required for efficient ubiquitination (Fig. 4a-c). To test whether direct mono-ubiquitination by BIRC6 followed the same requirements, we used recombinant RING domain (RING-R187A) and the RING domain lacking loop 2 (RING-ΔLoop2) as substrates in BIRC6-dependent ubiquitination assays. Consistent with cell-based results, the RING domain by itself was mono-ubiquitinated by BIRC6, while the RING mutant lacking the loop 2 region was not efficiently ubiquitinated (Fig. 5c). While full-length TRIM52 was also ubiquitinated on two additional minor sites, the RING domain in isolation was not (Fig. 5c), suggesting that indeed the major ubiquitination site for BIRC6 is inside the TRIM52 RING domain. Although our co-IP data indicated that BIRC6 binds outside of loop 2, we hypothesized that its strong negative charge may still contribute to its recognition. A positively charged CBD-3 domain arginine loop at the center of BIRC6 has been shown to mediate electrostatic interactions with a negatively charged region in two of its known substrates: HTRA2 and SMAC/DIABLO^52,53^. To test whether the same BIRC6 domain recognizes TRIM52, we used recombinant BIRC6 with a mutated CBD-3 arginine loop to negate its positive charge (3xRA)^52^. BIRC6-3xRA mono-ubiquitinated TRIM52 with a lower efficiency (Fig. 5d), indicating that the positive charge of the CBD-3 arginine loop region is partially required for the ubiquitination of TRIM52 by BIRC6.

BIRC6 mono-ubiquitinated TRIM52 *in vitro*, yet we identified abundant K48 poly-ubiquitin chains on TRIM52 isolated from cells (Fig. 2e-h, and Extended data table 3), suggesting that these are the major degradation signals on TRIM52. Therefore, we asked whether UBR4/KCMF1 and HUWE1 can extend the BIRC6-seeded mono-ubiquitination into poly-ubiquitin chains. To this end, we mono-ubiquitinated TRIM52 in the presence of BIRC6, immunoprecipitated TRIM52, and then used it as an input substrate in ubiquitination reactions with the indicated E3 ligases (Fig. 5e). In the presence of UBR4/KCMF1, high molecular weight poly-ubiquitin species appeared, consistent with transition into poly-ubiquitin chains (Fig. 5e-f). Together, these data indicate that TRIM52 is mono-ubiquitinated by BIRC6 on its RING domain, which can be extended by co-operation with UBR4/KCMF1 into poly-ubiquitin chains (Fig. 5g).

## Discussion

TRIM52 has been under positive selection pressure in humans and old-world primates. In contrast to other TRIM proteins with similar evolutionary patterns—like TRIM5α—no evidence has been found for direct anti-retroviral activity of TRIM52^6^, although some studies have reported a role in defenses against other viruses and inflammatory signaling^10,54–56^. In agreement with cell-intrinsic selection pressure, rather than pressure exclusively from exogenous pathogens, we found that *TRIM52* ablation decreased cellular fitness in a p53-dependent manner, in the absence of external stimuli^8,9^. This suggested a role of TRIM52 in maintaining primate genome integrity. Like *TRIM52*, many other regulators of the DNA damage response have been under positive selection pressure in humans and other primates^1,2^, although the selective advantages of these adaptations have remained poorly defined.

Here, we found that TRIM52 closely associates with TDP2 and ZNF451, known regulators of topoisomerase-dependent DNA lesions. Such lesions would be predominantly resolved through HDR and error-free NHEJ, yet can in part be compensated for by error-prone NHEJ^36,57,58^. Our genetic modifier screen showed that loss of key components of error-prone repair are synthetically lethal with loss of TRIM52, pointing to a physiologic role of TRIM52 in resolving topoisomerase lesions, and ultimate activation of HDR.

*TRIM52* ablation increased the concentrations of covalently-associated DNA-TOP2 complexes, indicating that TRIM52 is required for proper resolution of the irreversible TOP2-DNA tyrosyl cleavage complexes. Removal of the TOP2ccs from the DNA can be mediated through proteasomal degradation, or in a proteasome-independent manner^28–30,32^.

*TRIM52* arose by partial gene duplication of the evolutionary conserved *TRIM41* gene, which has been implicated in ubiquitination of TOP3B cleavage complexes and their subsequent proteasomal degradation^59^, suggesting that *TRIM52* divergent evolution may have resulted in an activity for TOP2. The precise stage and mechanism of TOP2cc resolution regulated by TRIM52, its cross-talk with TRIM41, and its effects on other topoisomerases will require additional future study.

RING E3 ligases interact with UBC folds in E2 conjugase enzymes, in part through interactions involving the two protruding RING loops^60^. Despite its disproportionately large and charge-biased loop 2, we unexpectedly found that TRIM52 had E3 ligase activity *in vitro* in combination with the promiscuous UBCH5B/C E2 conjugases. Especially since catalytic activity is not required for its own degradation^9^, it is thus tempting to speculate that TRIM52’s E3 activity may be required for its cellular function in the resolution of topoisomerase-mediated lesions.

Although we identified TRIM52 to be part of TDP2-containing complexes in cells, their individual ablation had different effects on TOP2cc levels, suggesting that these factors may function in parallel to each other. Consistent with previous findings in other cell models, *TDP2* knock-out alone had no measurable effect on TOP2cc levels on the DNA^25^, whereas these were increased in cells lacking *TRIM52*. In combination with TRIM52’s rapid proteasomal degradation, these findings suggest that TRIM52 may regulate resolution of TOP2 lesions. Although additional studies will be required to determine the relationship between TRIM52 degradation and its cellular function, we speculate that TRIM52 is either constantly degraded, yet stabilized when engaged in a DNA repair complex, or it is actively degraded when engaged in such complexes.

UBR4 operates with the UBE2A (RAD6) E2 conjugase enzyme^61^, which was identified alongside UBR4 and KCMF1 in our genetic screen. KCMF1 has been previously described as an interactor and co-factor of UBR4, although its function has remained enigmatic^46,62–65^. UBR4 was active *in vitro* in the absence of KCMF1, as it produced free ubiquitin chains, whereas KCMF1 by itself showed no E3 ligase activity. This suggests that KCMF1 could be a modular co-factor of UBR4, conferring recognition of mono-ubiquitinated substrates and facilitating E4 ubiquitin chain-extending activity to UBR4, or stabilizing the UBR4 protein itself. UBR4 and KCMF1 have been implemented in vesicular trafficking and autophagosomal/lysosomal protein degradation^46,62,66^, which our inhibitor studies indicate play no role in TRIM52 degradation. This raises the question how cells distinguish the ultimate destination of degradation for UBR4/KCMF1-dependent substrates.

The three giant E3 ligase components —HUWE1^48,67–73^, BIRC6^53,63,74–76^, and UBR4/KCMF1^46,66,77–84^— we identified to degrade TRIM52 have been previously implicated in the degradation of a wide range of substrates, thereby regulating various cellular processes. This has suggested that these E3 ligases can recognize multiple substrate classes based on wider biochemical or biophysical substrate features. Consistent with this notion, recent elucidation of the structure of HUWE1^49,50^ showed that this E3 ligase has three substrate binding modules allowing the recognition and ubiquitination of non-engaged nucleic acid-binding proteins and ubiquitinated/PARylated substrates. Likewise, BIRC6 binds and ubiquitinates various different apoptosis- and autophagy-related proteins^52,53,85^. Although such information is missing for UBR4 as its full-length structure has not been published to date, the fact that UBR4 has been found in independent screens to ubiquitinate aggregation-prone nascent polypeptides during proteotoxic stress^83^, various mitochondrial proteins^79,82^, and ER-associated degradation substrates^84^, indicates that also this giant E3 ligase may recognize various classes of substrates, potentially dependent on its association with different partners, such as KCMF1 and Calmodulin^46,86^.

Despite the fundamental insight that these E3s can recognize multiple substrate classes, *bona fide* cellular substrates and how they are recognized have remained sparse. This study has identified TRIM52 as a BIRC6 substrate for mono-ubiquitination. Our cellular mutagenesis data indicate that BIRC6 likely recognizes and binds the RING domain. This raises the question whether any of the other ∼600 RING domain proteins are likewise BIRC6 substrates. Moreover, BIRC6 had so far been shown to poly-ubiquitinate its substrates^52,85^, meaning it can both recognize substrates, modify them with the first ubiquitin, and extend this PTM into a poly-ubiquitin chain. While we cannot rule out that BIRC6 performs all of these actions on TRIM52 in cells in the presence of co-factors or PTMs, *in vitro* BIRC6 performed exclusively TRIM52 multi-mono-ubiquitination. This positions TRIM52 as an excellent substrate for future endeavors to elucidate how BIRC6 mono- and poly-ubiquitination activities are mechanistically controlled.

The fact that UBR4/KCMF1 and HUWE1 have been repeatedly identified to regulate partially overlapping substrates in cells could indicate the existence of a large protein complex consisting of some or all of the identified E3 ligases. In this context, cooperative action of BIRC6 and UBR4/KCMF1 controls activation of the Integrated Stress Response by ubiquitination of heme-regulated inhibitor (HRI)^63^. Therefore, like TRIM52, other cellular substrates may rely on sequential mono-ubiquitination by BIRC6, followed by UBR4/KCMF1 chain extension.

In summary, we identified a functional role of TRIM52 in DNA repair despite its non-conserved nature across mammals. These findings will enable future studies to identify what evolutionary benefit TRIM52 provides for humans, and perhaps brain complexity. In addition, we unraveled how one of the most unstable human proteins is degraded. We identified TRIM52 as a *bona fide* substrate for BIRC6 and UBR4/KCMF1, requiring its acidic loop 2 for recognition and degradation. While HUWE1 is functionally important for TRIM52 turn-over, its mechanistic targeting of TRIM52 will need further clarification in future studies. Together, these findings form the basis for follow-up studies addressing how TRIM52 turn-over is related to its cellular function.

## Acknowledgements

Next Generation Sequencing analysis Vienna Biocenter Core Facilities (VBCF) was performed by the Vienna Biocenter Core Facilities using the VBCF instrument pool. Proteomics analyses were performed by the Mass-spectrometry Facility at the Max Perutz Labs using the Max Perutz Labs and VBCF instrument pool; in particular we thank Markus Hartl and WeiQiang Chen for their expert support. Flow cytometry analyses were performed at the BioOptics FACS Facility at Max Perutz Labs using the Max Perutz Labs instrument pool; in particular we acknowledge Kitti Csalyi, Thomas Sauer, and Johanna Stranner for expert support. Microscopy was performed at the BioOptics Light Microscopy Facility at the Max Perutz Labs; we thank Thomas Peterbauer and Irmgard Fischer for their expert support and training. We are grateful to Robert Kurzbauer and Maria Heinke for purification of recombinant proteins. We thank Life Science Editors for editing services.

## Funding sources

This work was funded by Stand-Alone grants (P30231-B, P30415-B, P36572, P36945), Special Research Grant (SFB grant F79), and Doctoral School grant (DK grant W1261) from the Austrian Science Fund (FWF) to G.A.V., Austrian Science Fund Special Research Grant (FWF, SFB F 79) to T.C., and an ERC European Union’s Horizon 2020 research and innovation program grant (AdG 694978) to T.C. J.F.E. was supported by Austrian Science Fund DocFunds grant DOC 112-B. A.S. and V.B are recipients of the DOC fellowship of the Austrian Academy of Sciences. Research at the IMP is supported by Boehringer Ingelheim and the Austrian Research Promotion Agency (Headquarter grant FFG-852936).

## Materials availability statement

All data generated or analyzed during this study are included in the manuscript and supporting files.

## Funding Statement

The funders had no role in study design, data collection and interpretation, or the decision to submit the work for publication.

## Author contributions

Conceptualization, G.A.V., A.S., K.H., L.C., and T.C.; Methodology, G.A.V., A.S., K.H., J.F.E., V.B., A.M., and D.B.G.; Validation, A.S., K.H., J.F.E., and V.B; Formal Analysis, A.S., and K.H.; Investigation, A.S., K.H., J.F.E., V.B., A.M., and D.B.G.; Resources, J.F.E., and D.B.G.; Data Curation, A.S., K.H., and G.A.V.; Writing – Original Draft G.A.V., and A.S.; Writing – Review & Editing, A.S., K.H., J.F.E., V.B., A.M., D.B.G., L.C., T.C., and G.A.V.; Visualization A.S., and K.H.; Supervision G.A.V.; Project Administration G.A.V.; Funding Acquisition G.A.V.

## Declaration of interests

The authors declare no competing interests.

---

**Extended data figure 1-.**
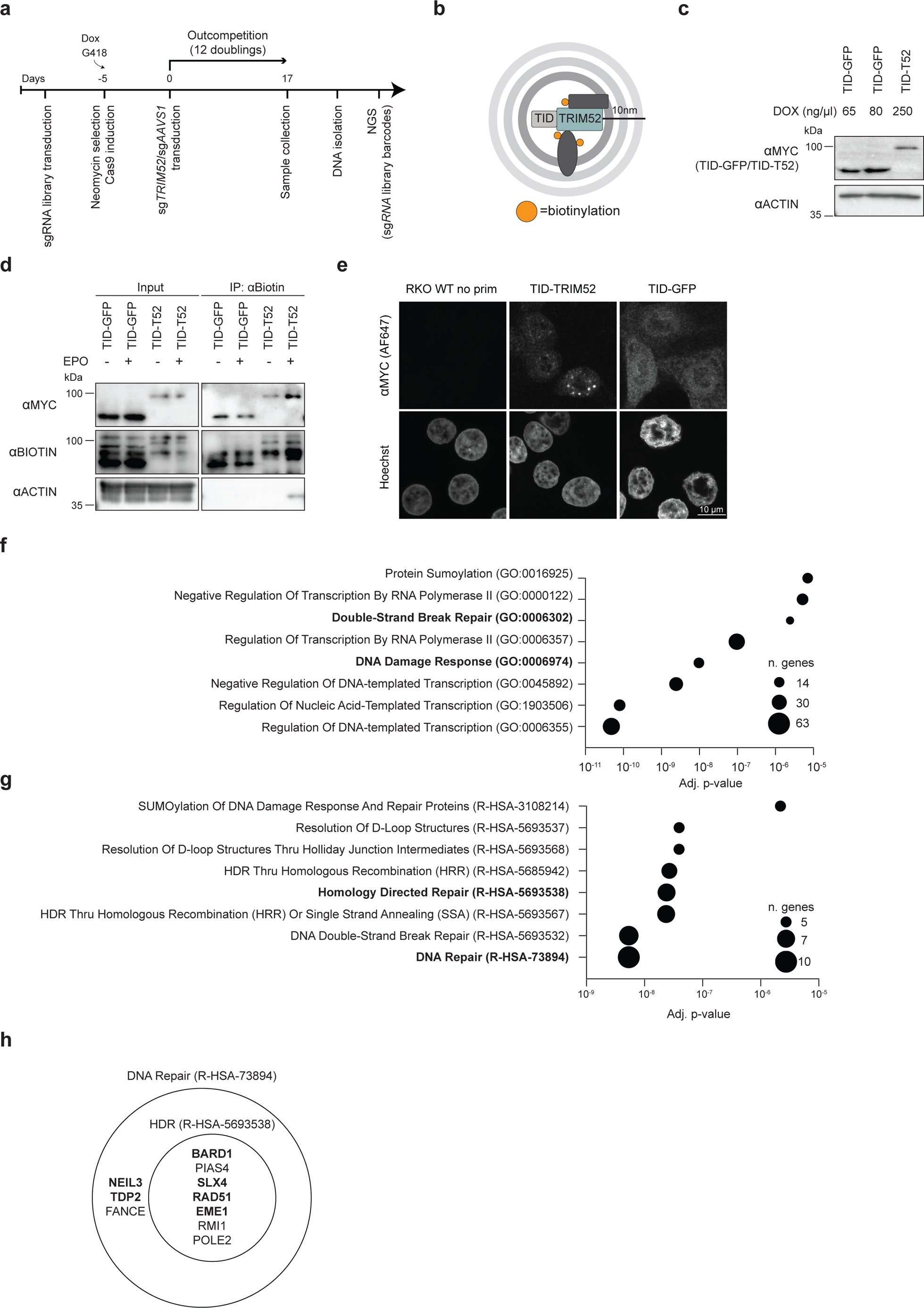
TRIM52 is required for resolution of TOP2 DNA damage lesions. (**a**) Timeline of the genetic modifier screen. (**b**) Schematic representation of TurboID proximity labelling principle. **(c)** RKO cells harboring Dox-inducible TurboID constructs were treated with the indicated concentrations of Dox, and protein expression analyzed by WB. (**d**) RKO cells expressing TurboID-TRIM52 or TurboID-EGFP fusion proteins were treated with epoxomicin for 5 h. and biotin for the last 15 min. Proteins were streptavidin affinity purified and analyzed by WB. (**e**) RKO cells stably expressing TurboID-TRIM52 and TurboID-EGFP were treated with epoxomicin for 5 h., fixed, and their subcellular localization determined by immunofluorescence confocal microscopy (scale bar: 10 µm). (**f**) Putative interactors of TRIM52 with p-value < 0.05 and Log2 fold change > 2.5 were selected and analyzed by gene ontology enrichment analysis. The enriched GO terms are plotted based on their adjusted p-value and the number of genes within each GO Biological Processes term. (**g**) Genes included in the DNA Damage Response (GO:0006974) and Double-strand Break Repair (GO:0006302) GO terms were further analyzed against the Reactome 2022 database. The enriched pathways are plotted based on their adjusted p-value and the number of genes within each Reactome 2022 term. (**h**) Venn-diagram describing the putative TRIM52 interactors in the DNA repair (R-HSA-73894) and HDR (R-HSA-5693538) GO terms.

**Extended data figure 2-.**
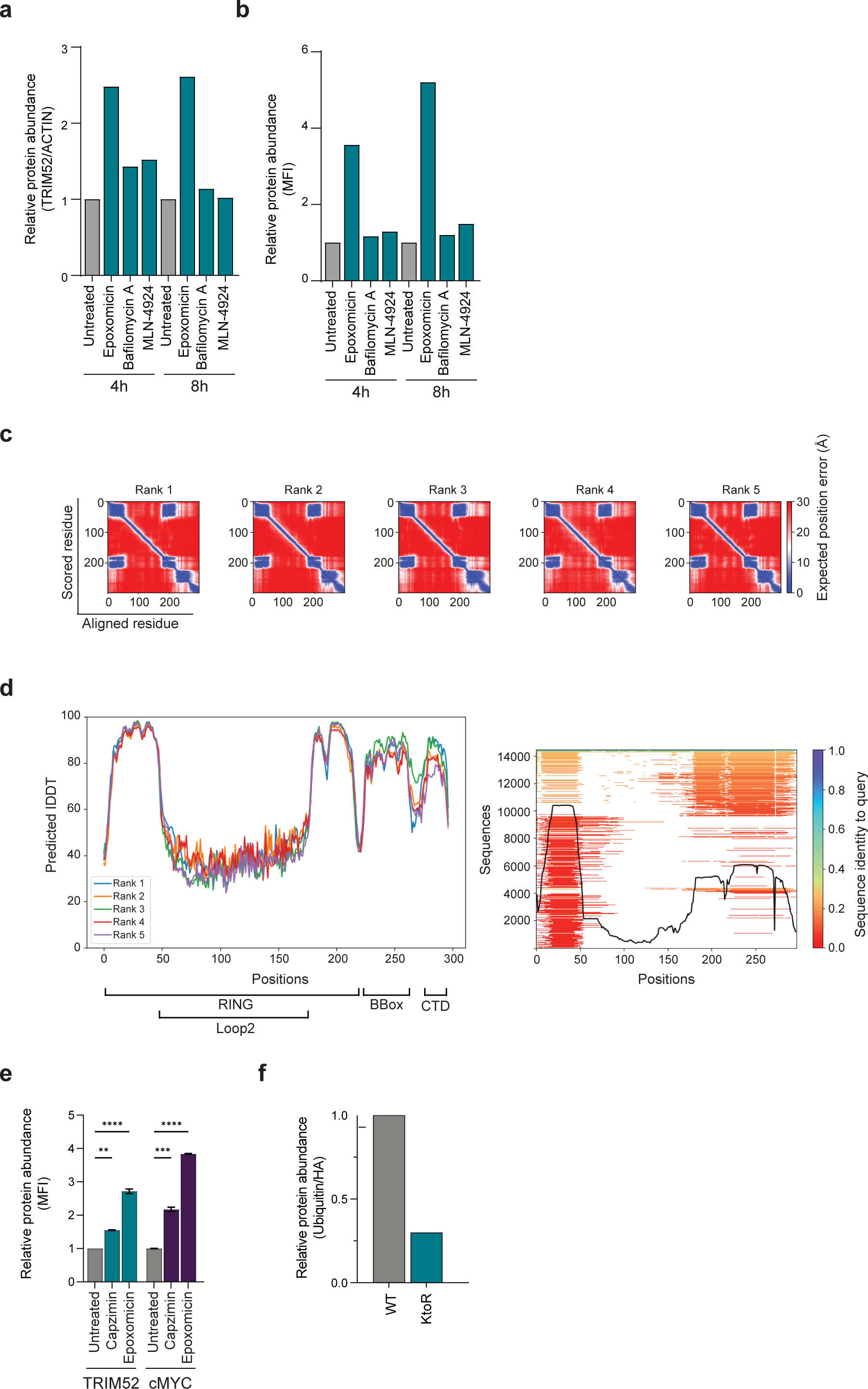
TRIM52 is degraded by the ubiquitin-proteasome system. (**a**) RKO cells expressing Ollas-tagged EGFP-TRIM52 were treated for 4 or 8 h. with epoxomicin, bafilomycin A, or MLN-4924. Ollas-EGFP-TRIM52 protein levels determined by WB, quantified and normalized to actin, or (**b**) by flow cytometry, their MFI quantified, normalized to MYC-mCherry and the untreated control and plotted. (**c**) Predicted aligned error plot of the AlphaFold2 model for TRIM52 structure prediction. (**d**) Predicted lDDT per position and sequence coverage of the AlphaFold2 model for TRIM52 structure prediction. (**e**) RKO cells expressing EGFP-TRIM52 or mCherry-cMYC fusion proteins were treated for 5 h. with the 20S inhibitor epoxomicin, or 19S-specific inhibitor capzimin. EGFP-TRIM52 and mCherry-cMYC protein levels were determined by flow cytometry, and MFI quantified. (Data represent biological samples, n = 2. Data were analyzed by 1-way ANOVA. ns: p > 0.05, *: p < 0.05, **: p < 0.01, ***: p < 0.001). (**f**) HEK-293T cells expressing HA-tagged WT TRIM52 or a lysine-to-arginine mutant were treated with epoxomicin for 5 h., after which TRIM52 was immunoprecipitated, and its ubiquitination analyzed by WB. The ubiquitin signal was quantified and normalized to WT HA-TRIM52 levels.

**Extended data figure 3-.**
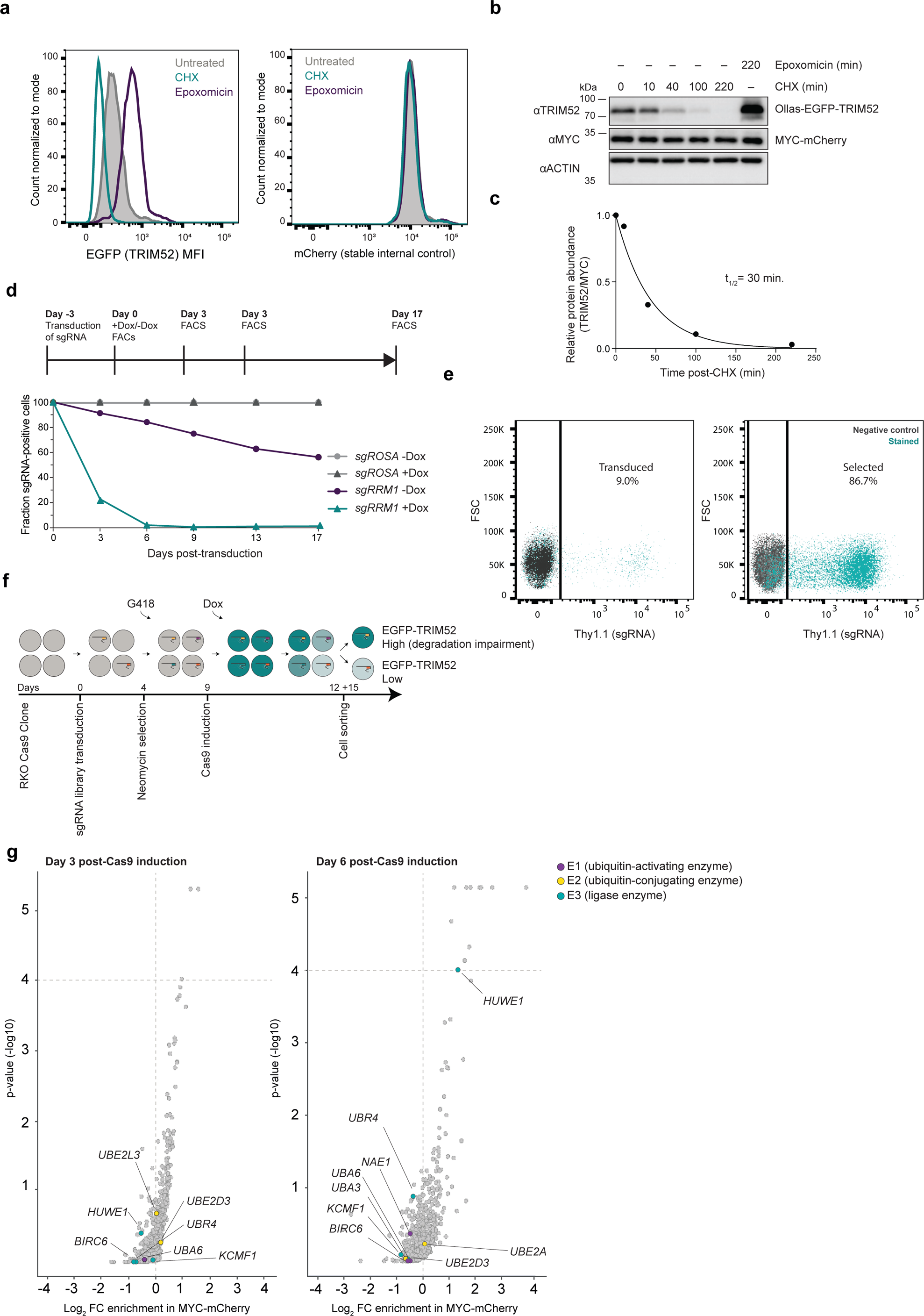
TRIM52 is targeted for degradation by multiple giant E3 ligases. (**a**) RKO cells were stably transduced with the TRIM52 reporter construct and single cell sorted based on mCherry and EGFP fluorescence intensities. Clones were expanded and treated for 4 h. with the proteasome inhibitor epoxomicin, or CHX. Ollas-EGFP-TRIM52 and MYC-mCherry protein levels were quantified by flow cytometry. (**b**) RKO cells stably transduced with the TRIM52 reporter construct were treated with epoxomicin or CHX for the indicated times. WCLs were analyzed by WB, (**c**) quantified and single-exponential decay curves were fitted to calculate half-life. (**d**) The screening cell line was transduced with sgRNAs targeting the essential gene *RRM1* or the *hROSA* safe harbor locus. Cells were mixed with WT cells and their cell fitness expressed as relative iRFP-positive fraction in the cell pool. (**e**) 4 days after transduction of the screening cell line with a lentiviral sgRNA library, the percentage of transduced cells was determined by analyzing the expression of Thy1.1 using flow cytometry. Transduced cells were selected using G418 for 4 days and Thy1.1 expression was analyzed using flow cytometry. (**f**) Schematic representation of FACS-based screen. (**g**) Enrichment plot of screen hits in the mCherry^high^ control sorted cell pool.

**Extended data figure 4-.**
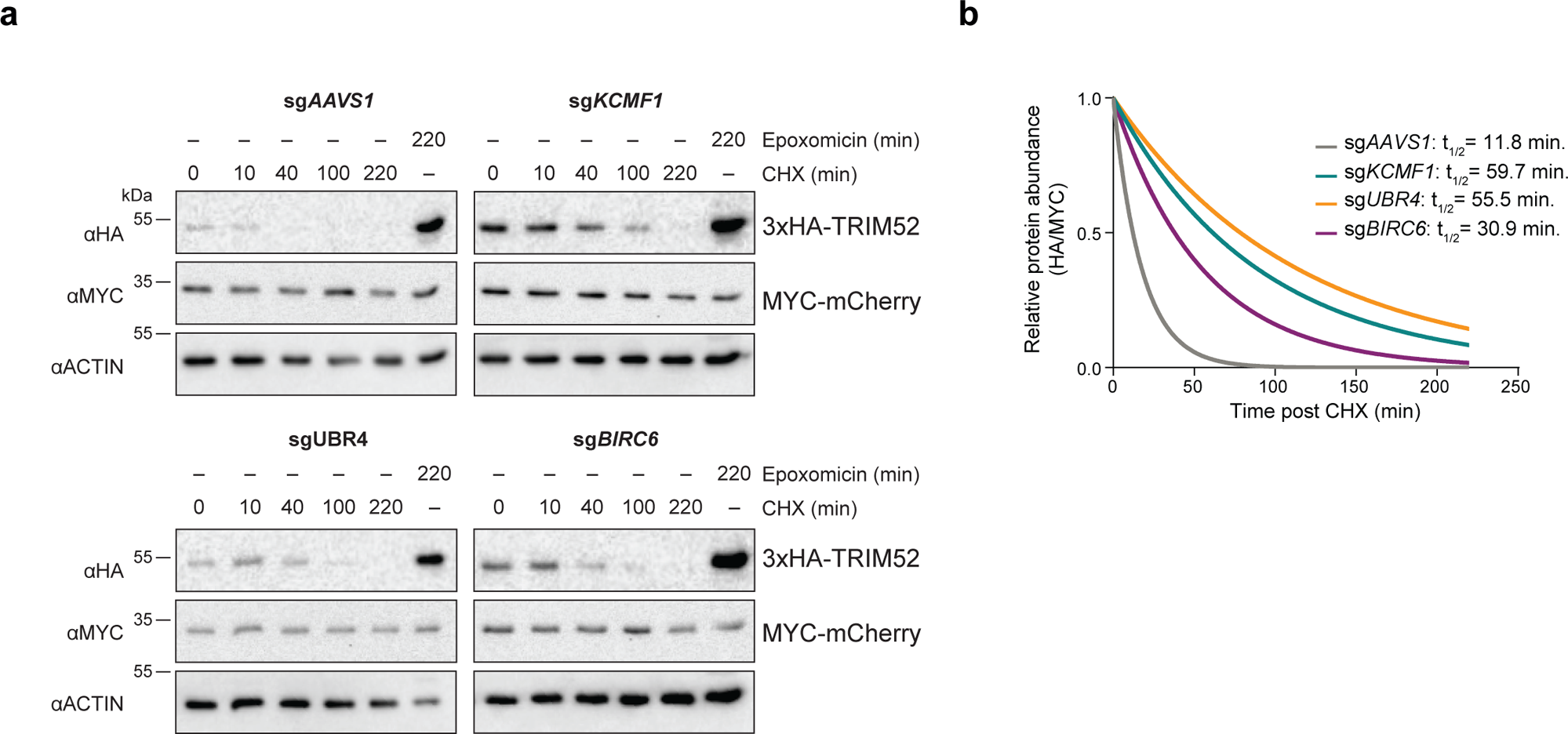
TRIM52 is targeted for degradation by multiple giant E3 ligases. (**a**) RKO cells stably expressing MYC-mCherry-P2A-HA-TRIM52 were treated with epoxomicin or CHX for the indicated times, after which WCLs were analyzed by WB, (**b**) quantified, and single-exponential decay curves were fitted to calculate half-lives.

**Extended data figure 5-.**
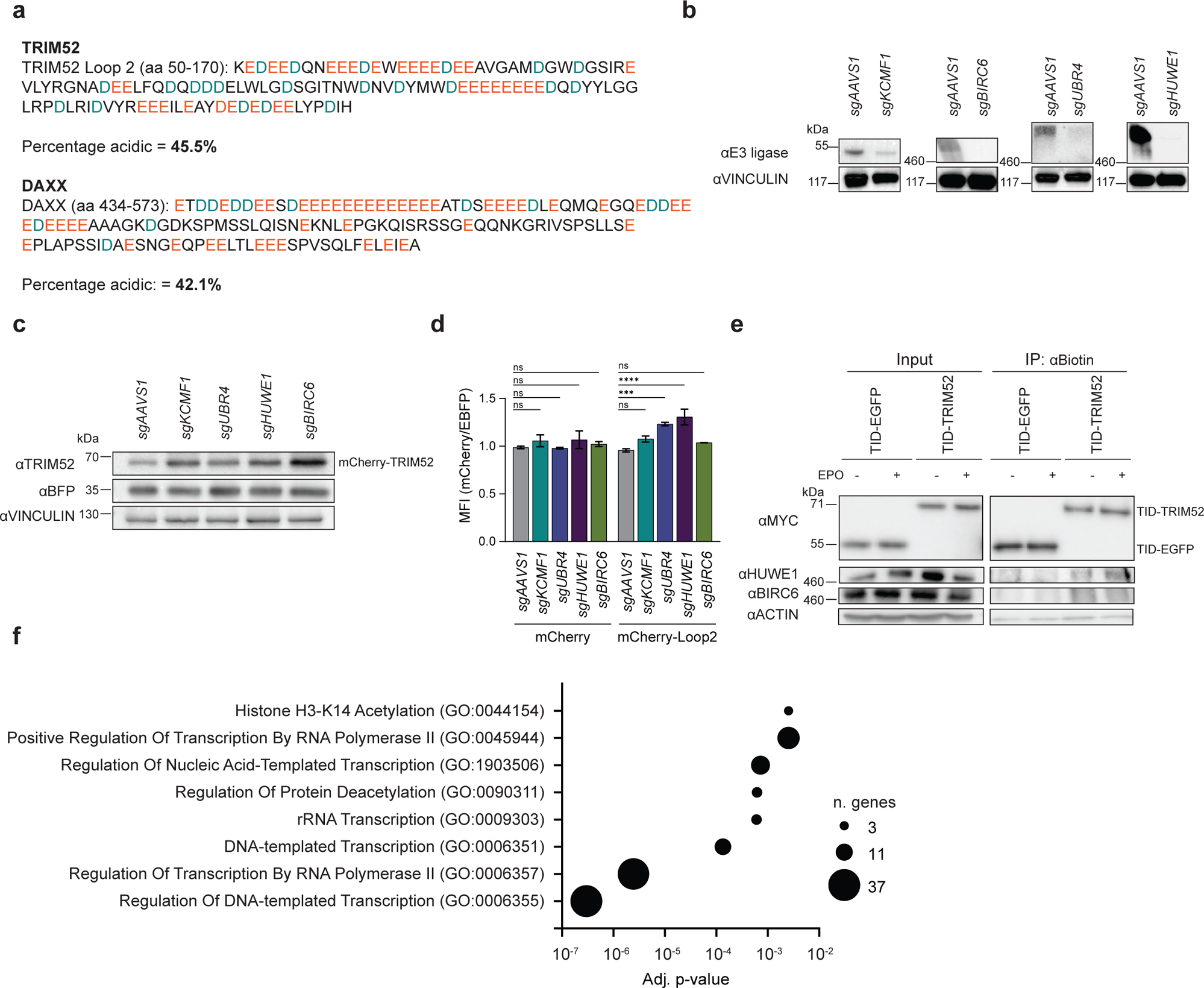
BIRC6, HUWE1, and UBR4/KCMF1 target the extended loop 2 region in the TRIM52 RING domain. (**a**) Sequences of acidic residue-rich regions in the loop 2 region of TRIM52 and DAXX. (**b**) HEK-293T cells constitutively expressing Cas9 and sgRNAs targeting the indicated genes were analyzed by WB. (**c**) HEK-293T constitutively expressing Cas9 and sgRNA targeting the indicated E3 ligases were transfected with plasmids expressing TwinStrep-mCherry-tagged TRIM52 and an internal EBFP control. Protein levels were analyzed by WB. (**d**) HEK-293T cells were transfected with plasmids expressing TwinStrep-mCherry or TwinStrep-mCherry-Loop2 and an internal EBFP control. Protein levels were measured by flow cytometry, normalized to EBFP and the sg*AAVS1* control and plotted (Data represent biological samples, n = 2. Data were analyzed by 2-way ANOVA. ns: p > 0.05, *: p < 0.05, **: p < 0.01, ***: p < 0.001, ****: p < 0.0001). (**e**) RKO cells expressing TurboID-TRIM52 or TurboID-EGFP fusion proteins were treated with biotin for 15 min., modified proteins were purified by streptavidin pull-down, and were analyzed by WB. (**f**) Putative interactors of TRIM52 with p-value < 0.05 and Log2 fold change > 2.5 were selected and analyzed by gene ontology enrichment analysis. The enriched terms are plotted based on their adjusted p-value and the number of genes within each GO Biological Processes term.

**Extended data figure 6-.**
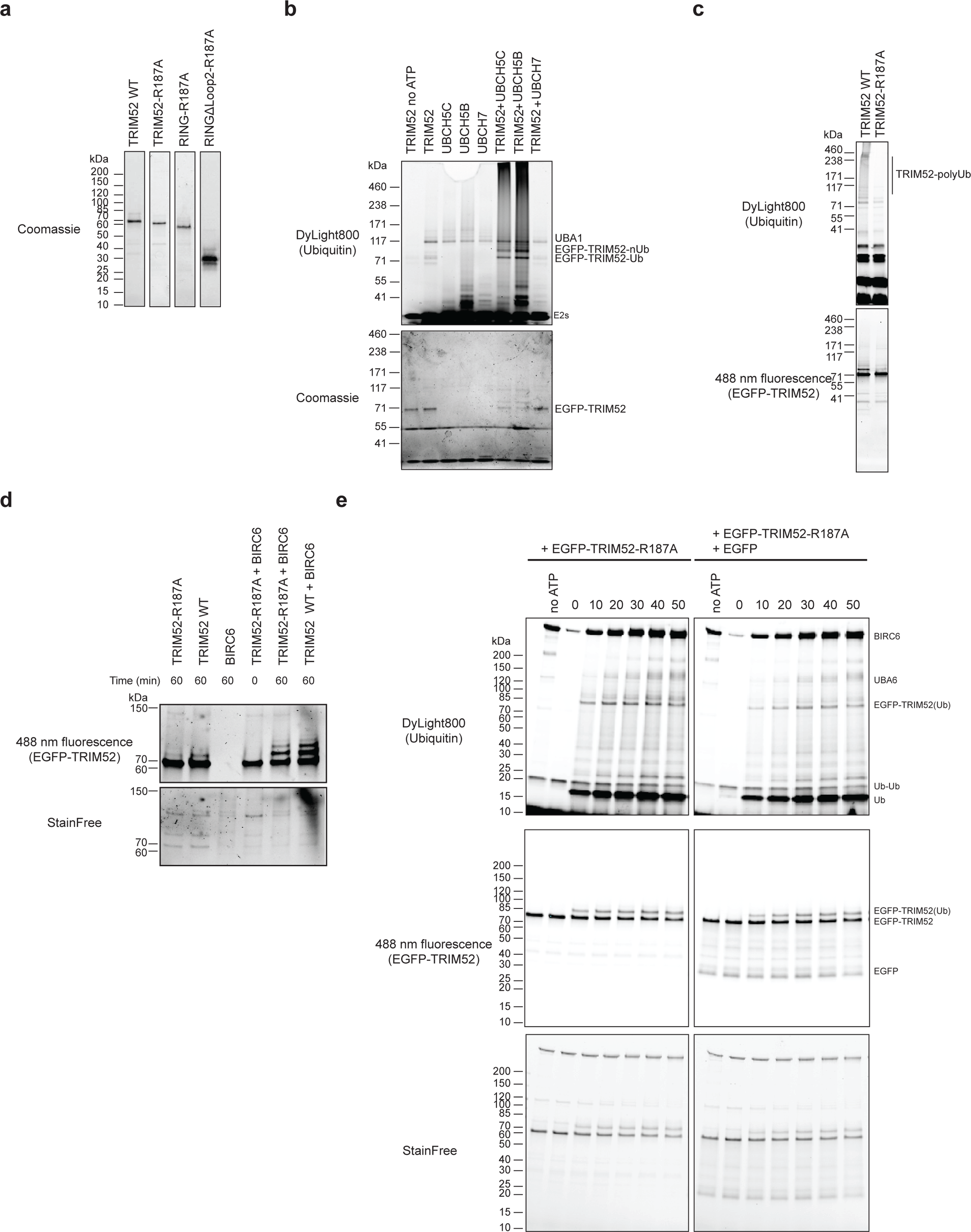
UBR4/KCMF1 extend poly-ubiquitin chains on sites directly mono-ubiquitinated by BIRC6. (**a**) TwinStrep-EGFP-TRIM52-WT, TwinStrep-EGFP-TRIM52-R187A, TwinStrep-EGFP-RING-R187A, TwinStrep-EGFP-RINGΔLoop2-R187A were expressed in Hi5 insect cells, recombinant protein purified by streptavidin affinity matrix and size-exclusion chromatography, and analyzed by SDS-PAGE with Coomassie staining. (**b**) Ubiquitination assay of EGFP-TRIM52-WT with different E2s, UBA1, and DyLight800-labeled ubiquitin. The reactions were incubated for 1h. at 37 °C in the presence of ATP. (**c**) Ubiquitination assay of WT EGFP-TRIM52 and its linchpin mutant (R187A) with UBCH5B, UBA1 and DyLight800-labeled ubiquitin. The reactions were incubated for 1h. at 37 °C in the presence of ATP. (**d**) Time-course ubiquitination assay of EGFP-TRIM52-WT and TRIM52-R187A with BIRC6, UBA6, and WT ubiquitin. The reactions were incubated for the indicated times at 37 °C in the presence of ATP. (**e**) Time-course ubiquitination assay of EGFP-TRIM52-R187A and EGFP as a control with BIRC6, UBA6, and DyLight800-labeled ubiquitin. The reactions were incubated for the indicated times at 37 °C in the presence of ATP.

## Materials and methods

### Cell culture

HEK-293T cells were cultured in high glucose Dulbecco’s Modified Eagle’s Medium (DMEM; Sigma-Aldrich, D6429) supplemented with 10% Fetal Bovine Serum (FCS; Sigma-Aldrich, F7524) and 1% Penicillin-Streptomycin (Sigma-Aldrich, P4333). RKO cells were cultured in Roswell Park Memorial Institute 1640 Medium (RPMI; Thermo Fisher Scientific, 21875) supplemented with 10% FCS (Sigma-Aldrich, F7524), 2% sodium pyruvate (Sigma-Aldrich, S8636), 1% Minimum Essential Medium (MEM) Non-Essential Amino Acids (Thermo Fisher Scientific, 11140050), and 1% Penicillin-Streptomycin (Sigma-Aldrich, P4333). All cells were cultured at 37 °C and 5% CO_2_ in a humidified incubator. Cells were treated with the following reagents for the indicated times: 200 μg/ml cycloheximide (CHX; Sigma-Aldrich, C1988); 10 μM MG132 (Sigma-Aldrich, M7449); 10 μM epoxomicin (Gentaur Molecular Products, 607-A2606); 200 ng/ml doxycycline hyclate (Dox; Sigma-Aldrich, D9891); 0.5-1 mg/ml G418 (Sigma-Aldrich, A1720); 400 nM bafilomycin A1 (Santa Cruz Biotechnology, sc-201550); 20 μM MLN-4924 (Abcam, ab216470), 5 μM etoposide (ETO, Sigma-Aldrich, E1383), 5 μM capzimin^40^. For gene targeting, HEK-293T cells were transduced with LentiCRISPR_V2 vectors (Addgene plasmid 52961; http://n2t.net/addgene:52961; RRID: Addgene_52961) encoding the indicated sgRNA sequences. Transduced cells were selected by supplementing DMEM culture media with 4 µg/ml puromycin (Invivogen, ant-pr-1). In RKO-Cas9 cells genome editing was induced using 200 ng/ml or 350 ng/ml final concentration of doxycycline hyclate (Dox, Sigma-Aldrich, D9891). Cas9 genome editing and expression in the absence of Dox from the TRE3G promoter was tested with competitive proliferation assays^8^. The cell lines, culture conditions and reagents used in this study are listed in the supplementary information. Cell lines used in this study were authenticated by STR analysis, and tested for mycoplasma contamination.

### Vectors

The lentiviral human sgRNA library was designed to targeted ubiquitin-proteasomal system-related genes, and is made up of 6 sgRNAs targeting each of the selected genes^19^. Lentiviral vectors expressing sgRNAs under a U6 promoter as well as selection colors EBFP or iRFP from a PGK promoter have been previously described^20^. sgRNA CDSs were cloned into pLentiv2-U6-PGK-iRFP670-P2A-Neo^20^ and used for gene targeting in RKO cell lines. The TRIM52 stability reporter (pLX-SFFV-MYC-mCherry-P2A-OLLAS-EGFP-TRIM52) was designed by cloning the open reading frame of human *TRIM52* into a modified pLX303 vector^47,87^. Single sgRNA CDSs were cloned in pLentiCRISPRv2 (Addgene plasmid 52961) to perform stable knock-outs in HEK-293T cells. cDNAs encoding TRIM52 FL, TRIM52 (KtoR), TRIM52 (2KtoR), TRIM52 (5KtoR), TRIM52 (ΔLoop2), RING, RING (ΔLoop2), RING (Loop2-neut), RING (Loop2-basic), DAXX (411-559), CY2B (1-103) were purchased from Twist Bioscience or generated by strand overlap PCR, and cloned into a modified pLX303 vector^87^. The plasmids and sgRNAs used in this study are listed in the supplementary information.

### Transfection of HEK-293T cells and production of virus-like particles

Transfection mixes were made containing DNA and polyethylenimine (PEI; Polysciences, 23966) in a ratio of 1:3 (μg DNA/μg PEI) in DMEM (Sigma-Aldrich, D6429) without supplements. The day prior to transfection, HEK-293T cells were seeded in 6-well clusters in supplemented DMEM media. For virus production, transfections were performed using 500 ng psPAX2 (Gag-Pol) plasmid (Addgene plasmid 12260; http://n2t.net/addgene:12260; RRID:Addgene_12260), 500 ng mini-genome plasmid and 100 ng pCMV2-VSVG plasmid^88^ in 6-well clusters. Transfected cells were incubated for 72 h. at 37 °C, after which virus-like particles were harvested by filtering the supernatant through a 0.45 μm filter. Virus-like particles were directly used after harvesting, or kept at 4 °C for short-term storage and at −80 °C for long-term storage.

### Cell competition assays

Competitive cell fitness assays were performed as described previously^89^. In brief, RKO cells harboring Dox-inducible Cas9 were transduced with iRFP or EBFP lentiviral sgRNA plasmids targeting the indicated genes. The multiplicity of transduction was such that 30-60% of cells were iRFP/EBFP-positive before the start of fitness measurements. Gene editing was induced with 200 ng/ml final concentration of Dox (Sigma-Aldrich, D9891) and the percentage of iRFP/EBFP-positive cells monitored for twenty days by flow cytometry at the indicated days. The relative fraction of sgRNA-positive cells was normalized to sg*hROSA or* sg*AAVS1* of the same cell line on day 0.

### CRISPR-iCas9 based modifier screens

RKO-iCas9-P2A-BFP cells carrying a genome-wide sgRNA library^19^ were grown in G418- (0.5 mg/ml, Sigma-Aldrich, A1720) and Dox-containing (200 ng/ml, Sigma-Aldrich, D9891) RPMI medium for iCas9 mediated base drop out of essential genes (T0). On day 5, cells were further transduced with VLPs for expression of a pair of sgRNAs targeting either *TRIM52*, *DGCR8* or the control locus *AAVS1* and selected in puromycin- (1 μg/ml) and Dox-supplemented RPMI. Cells were passaged for 12 doublings of control sgRNA (AAVS1) expressing cells (T3). All cells were grown at 500-fold library representation. The fraction of library positive cells was monitored regularly by Thy1.1 surface marker staining and flow cytometry. Samples for NGS sequencing were collected at timepoints T0 and T3. For harvesting, 1.5 x 10^8^ cells were pelleted, washed with PBS, and stored at −80°C until further processing.

### FACS-based CRISPR–iCas9 genetic screens

Lentivirus-like particles were used to transduce RKO-MYC-mCherry-P2A-OLLAS-EGFP-TRIM52 cells at a multiplicity of infection (MOI) of less than 0.2 TU/cell, and 1,000-fold library representation. The percentage of library-positive cells was determined after 4 days of transduction by immunostaining of the Thy1.1 surface marker and subsequent flow cytometric analysis. RKO cells with integrated lentiviral vectors were selected with G418 (1 mg/ml, Sigma-Aldrich, A1720) for 5 days, after which they were maintained in 0.5 mg/ml G418. After G418 selection, 20 million cells of unsorted reference samples were collected and stored at −80 °C until further processing. Cas9 genome editing was induced with Dox (200 ng/ml, Sigma-Aldrich, D9891) and after 3 and 6 days, cells were sorted by FACS. Cells were harvested, washed with PBS and sorted in fully supplemented RPMI-1640 using the FACS Aria III cell sorter operated by BD FACSDiva software (v8.0). RKO cells were gated for live, single, EBFP-positive (Cas9 expression), EGFP-positive, mCherry-positive, and. 2-3% of cells with the lowest and 2-4% of cells with the highest EGFP or mCherry signals were sorted into PBS. At least 1×10^6^ (EGFP^low^ and mCherry^low^) and 1×10^6^ (EGFP^high^ and mCherry^high^) cells were collected for each time point. Sorted samples were re-analyzed for purity, pelleted and stored at −80 °C until further processing. The gating strategy for flow cytometric cell sorting is shown in the Supplementary Information.

### Next-generation sequencing library preparation and genetic screen analysis

Next-generation sequencing (NGS) libraries of sorted and unsorted control samples were processed as previously described^20^. In brief, isolated genomic DNA was amplified with two-step PCR. The first PCR amplified the integrated sgRNA cassettes, and the second PCR introduced the Illumina adapters. Purified PCR products’ size distribution and concentrations were measured using a fragment analyzer (Advanced Analytical Technologies). Equimolar ratios of the obtained libraries were pooled and sequenced on a HiSeq 2500 platform (Illumina). Primers used for library amplification are listed in the supplementary information. Analysis of the CRISPR–Cas9 screen was performed as previously described^20^. In brief, sgRNAs enriched in day 3 and day 6 post-Cas9 induction-sorted samples were compared against the matching unsorted control populations harvested on the same days using MAGeCK^90^.

### RADAR assay

RADAR assays were performed as previously described^39^. In brief, 1 x 10^6^ RKO Cas9 cells were treated with etoposide (10 µg/ml) for 2 h., washed with PBS and lysed by adding 1 mL DNAzol (ThermoFisher Scientific, 11558626). Nucleic acids were precipitated by the addition of 0.5 mL 100% ethanol, incubation at −20 °C for 5 min., and centrifuged at 12,000 x g for 10 min. Precipitated nucleic acids were washed twice by addition of 75% ethanol and vortexing, resuspended in 200 μl TE buffer and heated at 65°C for 15 min. DNA was sheared by sonication (40% power for 15 s. pulse and 30 s. rest 5 times). Samples were centrifuged at 20,000 x g for 5 min. and supernatant-containing nucleic acids with covalently bound protein collected. Double-stranded DNA content of the sample was quantified by PicoGreen assay kit (Invitrogen, 11558626) and slot-blotted. TOP2cc were detected with anti-TOP2α antibody.

### Protein half-life determination

To estimate TRIM52 protein half-lives, RKO cell lines stably expressing MYC-mCherry-P2A-3xHA-TRIM52 or MYC-mCherry-P2A-3xHA-TRIM52-KtoR were treated with 200 μg/ml of cycloheximide (CHX, Sigma-Aldrich, C1988). At indicated time points, total protein extracts were generated using 1x disruption buffer (1.05 M Urea, 0.334 M β-Mercaptoethanol and 0.7% SDS) analyzed by WB, quantified, and normalized to stable internal control MYC-mCherry levels, and to time point 0 as indicated. Single exponential decay curves were plotted using GraphPad Prism (v9), from which protein half-lives were calculated.

### Co-immunoprecipitation assays

HEK-293T cells from one confluent 35-mm dish were lysed in 100 μl of Frackelton lysis buffer (10 mM Tris (pH 7.4), 50 mM NaCl, 30 mM Na_4_P_2_O_7_, 50 mM NaF, 2 mM EDTA, 1% Triton X-100, 1 mM DTT, 1 mM PMSF, and 1X protease inhibitor cocktail (cOmplete™ Protease Inhibitor Cocktail, 11697498001)). Cells were incubated on a rotating wheel at 4 °C for 30 min. then centrifuged at 20,000 x g at 4 °C for 30 min. Supernatants were transferred to new tubes and 10 μl (10% of the lysate) of each sample was collected as input fractions. Protein concentrations were determined by BCA Protein Assay Kit (Thermo Fisher Scientific, 23225) and 500 μg of lysates were incubated overnight at 4 °C on a rotating wheel with anti-HA antibody (Cell Signaling Technology, 1:100). The next day, magnetic beads (Protein A/G Magnetic Beads, Thermo Fisher Scientific, 88803), were blocked by rotation in 3% BSA in Frackelton Buffer for 1 h. at 4 °C. 25 μl of beads were added to 500 μg of lysates and rotated for 2 h. at 4 °C. Then, beads were washed five times with 1 ml of Frackelton buffer, and proteins eluted by boiling in 2X disruption buffer (2.1 M urea, 667 mM β-mercaptoethanol and 1.4% SDS) for 10 min. at 95 °C.

### Immunoprecipitations for ubiquitination

RKO cells stably expressing MYC-mCherry, MYC-mCherry-TRIM52 or MYC-mCherry-P2A-3xHA-TRIM52 were treated with 10 µM epoxomicin for 5 h. Cells from one confluent 35-mm dish were lysed in 100 µl of RIPA buffer with 1% SDS (50 mM Tris-HCl (pH 7.4), 150 mM NaCl, 1% SDS, 0.5% sodium deoxycholate, 1% Triton X-100), supplemented with 40 mM N-ethylmaleimide, 40 mM iodoacetamide, 25 U/ml benzonase, 1 mM PMSF, and 1X protease inhibitor cocktail (cOmplete™ Protease Inhibitor Cocktail, 11697498001). Cells were incubated on a rotating wheel at 4 °C for 30 min., and centrifuged at 20,000 x g at 4 °C for 15 min. Supernatants were transferred to new tubes. Protein concentrations were determined by BCA Protein Assay Kit (Thermo Fisher Scientific, 23225), and 30 µg of the lysates were collected as input. 500 μg of lysates were diluted 1:10 in RIPA buffer without SDS (50 mM Tris-HCl (pH 7.4), 150 mM NaCl, 0.5% sodium deoxycholate, 1% Triton X-100) and incubated with anti-HA antibody (Cell Signaling Technology, 1:100) overnight, or 25 µl of magnetic beads (RFP-Trap Dynabeads, Chromotek, rtd-20) for 2 h. Prior to incubation, beads were blocked by rotation in 3% BSA in RIPA Buffer for 1 h. at 4 °C. For the HA-IP, the next day 25 μl of magnetic beads (Protein A/G Magnetic Beads, Thermo Fisher Scientific, 88803) were added to 500 μg of lysates and rotated for 2 h. at 4 °C. Subsequently, beads were washed five times with 1 ml of RIPA buffer, supplemented with 600 or 300 mM NaCl respectively. Proteins were eluted by boiling in 2X disruption buffer (2.1 M urea, 667 mM β-mercaptoethanol and 1.4% SDS) for 10 min. at 95 °C.

### Immunoprecipitations for ubiquitination site identification by nLC-MS/MS

HEK 293T cells were transfected with MYC-mCherry-TRIM52 and 6xHis-Ubiquitin expression plasmids and after 48 h. treated with 10 µM epoxomicin (Gentaur Molecular Products, 607-A2606) for 5h. Cells from 5 confluent 15-cm dishes were lysed in 1 ml of RIPA buffer with 1% SDS (50 mM Tris-HCl (pH 7.4), 150 mM NaCl, 1% SDS, 0.5% sodium deoxycholate, 1% Triton X-100), supplemented with 40 mM N-ethylmaleimide, 40 mM iodoacetamide, 25 U/ml benzonase, 1 mM PMSF, and 1X protease inhibitor cocktail (cOmplete™ Protease Inhibitor Cocktail, 11697498001). Lysates were incubated on a rotating wheel at 4 °C for 30 min., and centrifuged at 20,000 x g at 4 °C for 30 min. Supernatants were transferred to new tubes. Protein concentrations were determined by BCA Protein Assay Kit (Thermo Fisher Scientific, 23225), and 30 µg of lysate was collected as input. The rest of the lysates were diluted 1:10 in RIPA no-SDS (50 mM Tris-HCl (pH 7.4), 150 mM NaCl, 0.5% sodium deoxycholate, 1% Triton X-100) and incubated with 250 µl of magnetic beads (RFP-Trap Dynabeads, Chromotek, rtd-20) for 2 h. Subsequently, beads were washed five times with 1 ml of RIPA wash buffer (50 mM Tris-HCl (pH 7.4), 600 mM NaCl, 0.1% SDS, 0.5% sodium deoxycholate, 1% Triton X-100) supplemented with 600 mM NaCl. Protein was eluted from the beads in 1 ml elution buffer (50 mM Tris-HCl (pH 8), 150 mM NaCl, 0.1% SDS, 4M urea, 5mM imidazole), supplemented with 40 mM N-ethylmaleimide, 40 mM iodoacetamide. Samples were then diluted 1:10 in wash buffer (50 mM Tris-HCl (pH 8), 150mM NaCl, 4 M urea, 5 mM imidazole) and incubated with 500 µl Ni-NTA beads (Thermo Fisher Scientific, 88831) at 4 °C for 2 h. Beads were washed 5 times in 1 ml of wash buffer on a rotating wheel at 4 °C for 5 min. After washes, beads were re-suspended in 100 µl 50 mM Tris-HCl (pH 8) and analyzed by nLC-MS/MS.

### TurboID proximity labelling

The TurboID sequence from plasmid V5-TurboID-NES_pCDNA3^91^ (Addgene plasmid: 107169) was PCR amplified and integrated into a modified version of pCW57.1^92^. TRIM52 and EGFP were cloned into this plasmid for their expression with an N-terminal TurboID tag. Subsequently, RKO cell lines stably expressing various Dox-inducible TurboID constructs were created. For the TurboID experiments cell lines expressing Dox-inducible TurboID-TRIM52 and TurboID-EGFP were stimulated with 55 or 250 ng/ml doxycycline hyclate (Dox; Sigma-Aldrich, D9891) for 48 h. to induce the expression of the TurboID fusion constructs. Subsequently, cells were stimulated for 4 h. with 10 µM epoxomicin (Gentaur Molecular Products, 607-A2606) to inhibit proteasomal degradation where indicated or left untreated, and finally biotinylation was induced for 15 min. by addition of 500 µM biotin (Sigma, B4501) to the cell culture medium. Biotinylated proteins were enriched with streptavidin-coated beads. In brief, cells were washed four times with ice-cold PBS, prior to lysis in RIPA lysis buffer (50 mM Tris HCl (pH 7.4), 150 mM NaCl, 1% NP-40, 0.5% Sodium Deoxycholate, 1 mM EDTA, 0.1% SDS, 1 mM PMSF and 1X protease inhibitor (cOmplete™ Protease Inhibitor Cocktail, 11697498001)). Lysates were rotated for 15 min. at 4 °C, centrifuged at 18,500 x g for 10 min. at 4 °C, and protein concentrations were determined by BCA assay. 1200 µg of protein were incubated overnight rotating at 4°C with 200 µl of streptavidin beads (Thermo Scientific, 88816) which were acetylated with Sulfo-NHS-Acetate beforehand, as described^93^. Beads were washed twice with 1 ml of RIPA buffer, once with 1 ml of 2 M Urea in 10 mM Tris (pH 8), twice with 1 ml RIPA buffer and five times with 50 mM HEPES (pH 7.8). Three technical replicates were subjected to nLC-MS/MS analysis.

For nLC-MS/MS sample preparation, the beads were resuspended in 40 µl 1 M urea and 50 mM ammonium bicarbonate. Disulfide bonds were reduced with 1.6 µl of 250 mM dithiothreitol (DTT) for 30 min. at RT before adding 1.6 µl of 500 mM iodoacetamide and incubating for 30 min. at RT in the dark. Remaining iodoacetamide was quenched with 0.8 µl of 250 mM DTT for 10 min. Proteins were digested with 150 ng LysC (mass spectrometry grade, FUJIFILM Wako chemicals) at 25°C overnight. The supernatant was transferred to a new tube and digested with 150 ng trypsin (Trypsin Gold, Promega) in 1.5 µl 50 mM ammonium bicarbonate at 37°C for 5 h. The digest was stopped by the addition of trifluoroacetic acid (TFA) to a final concentration of 0.5 %, and the peptides were desalted using C18 Stagetips^94^.

### Co-immunoprecipitation of purified proteins

1 µg of purified EGFP-TRIM52 (WT or domain mutants) and 1 µg of purified HUWE1 were incubated in ubiquitination buffer (25 mM HEPES (pH 7.5), 150 mM KCl, 4 mM MgCl_2_, 0.5 mM TCEP) supplemented with 1x protease inhibitor cocktail (cOmplete™ Protease Inhibitor Cocktail, 11697498001) for 2 h. at 4 °C on a rotating wheel. In the meantime, GFP-Trap magnetic beads (Chromotek, gtd-20) were blocked by rotation in 3% BSA ubiquitination buffer for 1 h. at 4 °C. After the incubation, 5% of the reaction volume was aliquoted for the input fraction. The rest of the sample was incubated with 10 µl magnetic beads for 30 min. at 4 °C on a rotating wheel. Beads were washed 5 times with 500 µl ubiquitination reaction buffer. Proteins were eluted by boiling in 2X disruption buffer (2.1 M urea, 667 mM β-mercaptoethanol and 1.4% SDS) for 10 min. at 95 °C and analyzed by SDS-PAGE and WB.

### *In vitro* ubiquitination assays

*In vitro* ubiquitination assays contained 20 μM ubiquitin (WT, K0 or FLAG-tagged), 1 μM DyLight800/FLAG-labeled ubiquitin for in-gel visualization, 0.2 μM E1 (UBA6, UBA1), 0.4 μM E2 (RAD6, UBCH5B) 0.4 μM E3 (BIRC6, UBR4, HUWE1, KCMF1) and 2 μM substrate protein (EGFP-TRIM52), unless indicated otherwise. The assays were performed in 25 mM HEPES pH 7.5, 150 mM KCl, 4 mM MgCl_2_, 0.5 mM TCEP (assay buffer) at 37 °C for the indicated times in 10-20 μl volumes. Reactions were initiated by the addition of 5 mM ATP and stopped by adding 2x reducing SDS-PAGE loading buffer. SDS-PAGE was performed using 4–20% Mini-PROTEAN TGX Stain-Free (BioRad, 4568094) gels. A Bio-Rad ChemiDoc MP system was used for in gel fluorescence imaging.

### Immunofluorescence confocal microscopy

RKO cells stably expressing TurboID-TRIM52 and TurboID-EGFP were treated with Dox (1 μg/ml) for 4 days to induce TurboID expression, seeded onto coverslips. After 48h., cells were treated with 10 μM epoxomicin (Gentaur Molecular Products, 607-A2606) for 5 h. Cells were washed once with PBS, fixed on coverslips with 4% PFA for 15 min. at RT and washed twice with PBS for 5 min. Cells were permeabilized with PBS-0.25% Triton X-100 for 5 min. at RT and washed three times with PBS before blocking for 30 min. in blocking solution (PBS, 1% BSA). Cells were stained with anti-MYC antibody (4A6, 1:500 in 1% BSA) for 1 h. at 37°C, in a moisture chamber, washed three times with PBS for and subsequently stained with AF647 anti-mouse secondary antibody (1:200, Abcam, ab169348 in 1% BSA) for 1 h at RT, protected from light. Cells were washed three times with PBS and incubated with 0.4X Hoechst (Thermo Fisher Scientific) in PBS. The coverslips were mounted using ProLong™ Gold Antifade Mountant (Invitrogen). Images were collected using a Zeiss LSM 980 confocal microscope at 40X magnification.

### Western blot analysis

Cells were lysed in 1x disruption buffer or RIPA lysis buffer supplemented with 1% SDS and 25 U/ml benzonase. Lysates were rotated for 30 min. at 4 °C and then centrifuged at 18,500 x g for 10 min. at 4 °C. Supernatants were transferred to new tubes and protein concentrations were determined by BCA Protein Assay Kit (Thermo Fisher Scientific, 23225). Between 20-50 μg of protein per sample was mixed with Laemmli sample buffer (62.5 mM Tris-HCl (pH 6.8), 5.8% glycerol, 2% SDS and 1.7% β-mercaptoethanol), and boiled for 10 min. Proteins were loaded on 10% SDS polyacrylamide gels, or alternatively 4–20% Mini-PROTEAN TGX Stain-Free (BioRad, 4568094) or 3-8% NuPage gels (Invitrogen, EA03785BOX) to probe for high molecular weight proteins. Proteins were separated by SDS-PAGE using Tris-Glycine (25 mM Tris, 192 mM glycine, 0.1% SDS) or Tris-Acetate (2.5 mM Tricine, 2.5 mM Tris, 0.05% SDS) SDS running buffer, respectively. Proteins were blotted on PVDF membranes at 4 °C for 1 h and 15 min. at 300 mA in Towbin buffer (25 mM Tris pH 8.3, 192 mM glycine and 20% ethanol). Membranes were blocked in 5% BSA in PBS-T for 1 h. at RT, and subsequently incubated with primary antibodies diluted in 5% BSA overnight at 4 °C. The next day, membranes were washed three times for 5 min. each with PBS-0.05%-Tween20 and incubated with HRP-coupled secondary antibodies in 5% skimmed milk for 1 h. at RT and imaged with the ChemiDoc Imaging System (Bio-Rad). Relative protein levels were quantified using Image Lab software (Bio-Rad). The antibodies used in this study are listed in Table S6.

### Sample preparation for nLC-MS/MS analysis

Beads were resuspended in 96 µl 50 mM ammonium bicarbonate and reduced with 2 µl of 50mM TCEP for 30 min. at RT before adding 2 µl of 200 mM MMTS and incubating for 30 min. at RT in the dark. Afterwards, samples were digested with 300 ng trypsin (Trypsin Gold, Promega; V5280) at 37 °C overnight. Supernatants were transferred to new tubes and digests were stopped by the addition of trifluoroacetic acid (TFA) to a final concentration of 0.5 %, and peptides were desalted using C18 Stagetips^94^. Half of the trypsin digests were dried in a SpeedVac and resuspended in 100 mM Tris-HCl (pH 8.5). Samples were further digested with 50 ng chymotrypsin at 25°C for 5 h. The digests were stopped by the addition of trifluoroacetic acid (TFA) to a final concentration of 0.5 %, and the peptides were desalted using C18 Stagetips^94^.

### Liquid chromatography-mass spectrometry data acquisition and analysis

For the identification of ubiquitinated residues, peptides were separated on an Ultimate 3000 RSLC nano-flow chromatography system (Thermo Fisher Scientific), using a pre-column for sample loading (Acclaim PepMap C18, 2 cm × 0.1 mm, 5 μm, Thermo Fisher Scientific), and a C18 analytical column (Acclaim PepMap C18, 50 cm × 0.75 mm, 2 μm, Thermo Fisher Scientific), applying a segmented linear gradient from 2% to 35% and finally 80% solvent B (80 % acetonitrile, 0.1 % formic acid; solvent A 0.1 % formic acid) at a flow rate of 230 nl/min over 60 min. Eluting peptides were analyzed on an Exploris 480 Orbitrap mass spectrometer (Thermo Fisher Scientific) coupled to the column with a FAIMS pro ion-source (Thermo Fisher Scientific) using coated emitter tips (PepSep, MSWil) with the following settings: the mass spectrometer was operated in DDA mode with two FAIMS compensation voltages (CV) set to −35, −45, −60 or −75 and 0.8 s cycle time per CV. The survey scans were obtained in a mass range of 350-1500 m/z, at a resolution of 60k at 200 m/z, and a normalized AGC target at 100%. The most intense ions were selected with an isolation width of 1.2 m/z, fragmented in the HCD cell at 28% collision energy, and the spectra recorded for max. 100 ms at a normalized AGC target of 200% and a resolution of 30k. Peptides with a charge of +2 to +6 were included for fragmentation, the peptide match feature was set to preferred, the exclude isotope feature was enabled, and selected precursors were dynamically excluded from repeated sampling for 20 seconds. MS raw data split for each CV using FreeStyle 1.7 (Thermo Fisher Scientific), were analyzed using the MaxQuant software package (version 2.1.0.0)^95^ with the Uniprot human reference proteome (version 2022_01, www.uniprot.org), target protein sequences, as well as a database of most common contaminants. The search was performed with trypsin/chymotrypsin specificity and a maximum of two or four missed cleavages at a protein and peptide spectrum match false discovery rate of 1%. MMTS of cysteine residues was set as fixed, GlyGly(K), oxidation of methionine, and N-terminal acetylation as variable modifications - all other parameters were left at default. The mass spectrometry proteomics data have been deposited to the ProteomeXchange Consortium via the PRIDE partner repository^96^ with the dataset identifier PXD051295.

For TurboID proximity labelling analysis, peptides were separated on an Ultimate 3000 RSLC Nano-flow chromatography system (Thermo Fisher Scientific), using a pre-column for sample loading (Acclaim PepMap C18, 2 cm × 0.1 mm, 5 μm, Thermo Fisher Scientific), and a C18 analytical column (Acclaim PepMap C18, 50 cm × 0.75 mm, 2 μm, Thermo Fisher Scientific), applying a segmented linear gradient from 2% to 35% and finally 80% solvent B (80 % acetonitrile, 0.1 % formic acid; solvent A 0.1 % formic acid) at a flow rate of 230 nl/min over 120 min. Eluting peptides were analyzed on an Exploris 480 Orbitrap mass spectrometer (Thermo Fisher Scientific) coupled to the column with a FAIMS pro ion-source (Thermo Fisher Scientific) using coated emitter tips (PepSep, MSWil) with the following settings: the mass spectrometer was operated in DDA mode with two FAIMS compensation voltages (CV) set to −45 or −60 and 1.5 s cycle time per CV. The survey scans were obtained in a mass range of 350-1500 m/z, at a resolution of 60k at 200 m/z, and a normalized AGC target at 100%. The most intense ions were selected with an isolation width of 1 m/z, fragmented in the HCD cell at 28% collision energy, and the spectra recorded for max. 50 ms at a normalized AGC target of 100% and a resolution of 15k. Peptides with a charge of +2 to +6 were included for fragmentation, the peptide match feature was set to preferred, the exclude isotope feature was enabled, and selected precursors were dynamically excluded from repeated sampling for 45 seconds. MS raw data split for each CV using FreeStyle 1.7 (Thermo Fisher Scientific), were analyzed using the MaxQuant software package (version 1.6.17.0)^95^ with the Uniprot human reference proteome (version 2021_03, www.uniprot.org), as well as a database of most common contaminants. The search was performed with full trypsin specificity and a maximum of two missed cleavages at a protein and peptide spectrum match false discovery rate of 1%. Carbamidomethylation of cysteine residues was set as fixed, oxidation of methionine, phosphorylation (STY) and N-terminal acetylation as variable modifications. For label-free quantification the “match between runs” only within the sample batch and the LFQ function were activated - all other parameters were left at default. MaxQuant output tables were further processed in R 4.2.1 (https://www.R-project.org) using Cassiopeia_LFQ (https://github.com/moritzmadern/Cassiopeia_LFQ). Reverse database identifications, contaminant proteins, protein groups identified only by a modified peptide, protein groups with less than two quantitative values in one experimental group, and protein groups with less than 2 razor peptides were removed for further analysis. Missing values were replaced by randomly drawing data points from a normal distribution model on the whole dataset (data mean shifted by −1.8 standard deviations, a width of the distribution of 0.3 standard deviations). The mass spectrometry proteomics data have been deposited to the ProteomeXchange Consortium via the PRIDE partner repository with the dataset identifier PXD051272.

### Protein purification

Baculovirus encoding the respective TwinStrep tagged EGFP-TRIM52 constructs (TRIM52-WT, TRIM52-R187A, RING-R187A, RING ΔLoop2) were added to 500 mL of 1×10^6^ mL^−1^ Trichoplusia ni High Five (Accession: CVCL_C190) insect cells at a ratio of 1:100, grown in ESF921 serum-free media. Cell growth continued at 27 °C at 100 rpm for three days, after which cells were pelleted at 500 x g for 10 min. and resuspended in 25 mM HEPES pH 7.5, 300 mM KCl before flash freezing. For purification, cells were thawed and mixed with two mini c0mplete EDTA free protease inhibitor tablets (Roche) and bezonase (Molecular Biology Services, IMP). Cells were dounced 10 times with a glass douncer leading to mechanical lysis, and supernatant was collected after a 45 min, centrifugation step at 19000 rpm. The sample was applied to a 1 mL StrepTrap column (Cytiva), washed, and eluted in the same buffer with 3 mM d-desthiobiotin. Peak fractions were pooled, concentrated, and loaded onto a Superdex 200 10/300 (Cytiva) column in 25 mM HEPES pH 7.5, 300 mM KCl via a 1 mL loop. Fractions and purity were chosen and assessed by SDS-PAGE and the resulting protein was concentrated and frozen.

Recombinant human BIRC6, UBCH5B, UBA1, UBA6, Ubiquitin, DyLight800-labeled Ubiquitin were purified as previously described^52,85^. Human HUWE1 was codon-optimized, synthesized in fragments and assembled into a GoldenBac pGBdest vector via a BsaI-GoldenGate reaction cloning^97^. The construct contains a His8-tag, a rigid enhancer linker (AEAAAKEAAAKEAAAKEAAAKALEAEAAAKEAAAKEAAAKEAAAKA) followed by a TEV cleavage site. Human UBR4 was cloned by the same approach, but instead with a C-terminal Strep tag which is separated from the protein via a 3C cleavage site. Expression plasmids were transformed into the DH10 MultiBac cells for bacmid generation. *Spodoptera frugiperda* (Sf9) cells were cultured in ESF921 serum-free growth medium (Expression Systems) and transfected with the bacmids for virus production, which was quantified using yellow fluorescent protein signal. HUWE1 and UBR4 were then expressed in *Trichoplusia ni* High-Five insect cells (Thermo Fisher Scientific) at a density of 1.5 x10^6^. The cells were inoculated with a 1:70 dilution from the V1 stock for 92 h. at 21°C in Insect Xpress Protein-free Insect Cell Medium (Lonza) supplemented with GlutaMAX (GIBCO) and Pen/Strep amphotericin B (Lonza). Cells were harvested by centrifugation at 700 x g, washed in PBS, flash-frozen and stored at −70°C.

For HUWE1 purification, the cell pellet was thawed and resuspended in 50 mM HEPES pH 7.5, 300 mM NaCl, 0.5 mM TCEP, 20 mM imidazole, supplemented with Complete EDTA-free Protease inhibitor (Roche) and 20 µl benzonase (IMP Molecular Biology Service) and lysed using a douncer. The supernatant was separated by centrifugation at 40,000 x g. and the soluble fraction loaded on a 5 ml HisTrap HP column (Cytiva). The column was pre-equilibrated in the lysis buffer using an Äkta Pure 25 system (Cytiva). The column was washed with 10 CVs of the same buffer, followed by 7 CVs of buffer supplemented with 35 mM imidazole, and then eluted with 300 mM imidazole. The protein was cleaved with TEV protease and simultaneously dialyzed overnight to remove the imidazole. The cleaved protein was then reapplied to HisTrap HP column in 20 mM imidazole and flow through collected and concentrated to 1.5 ml. Following that, the protein was applied to a Superose 6 16/60 column (Cytiva) equilibrated in 50 mM HEPES pH 7.5, 150 mM NaCl, 0.5 mM TCEP and HUWE1-containing fractions were pooled and concentrated. The protein purity was assessed by SDS-PAGE and the concentration was estimated by absorption at A280 nm using an extinction coefficient of 251,770 M^−1^ cm^−1^.

For UBR4 purification, the same lysis procedure was used except with phosphate buffered saline (PBS), and 0.5 mM TCEP at pH 7.4 as the buffer. The soluble lysate was loaded on a 5 ml StrepTrap HP column (Cytiva) equilibrated in PBS, washed with the same buffer and eluted with 2.5 mM desthiobiotin. Eluted UBR4 was then applied to a Resource Q column (Cytiva) equilibrated in PBS for additional purification via anion exchange chromatography using a 250 to 500 mM NaCl gradient. UBR4-containing fractions were pooled and concentrated. The concentration was measured by absorption at 280 nm using a calculated extinction coefficient of 472,140 M^−1^cm^−1^.

The codon-optimized UBE2A (RAD6) gene fragment was synthesized by Twist Bioscience. The NEB HiFi assembly kit (New England Biolabs) was used to clone the gene fragment into a pET29b expression vector, which also contained an N-terminal His6-tag and TEV cleavage site. The protein was expressed from *E. coli* BL21 (DE3) cells using IPTG induction at 20 °C overnight. Cells were resuspended in buffer containing 50 mM Tris (pH 7.5), 150 mM NaCl, 10 mM imidazole, supplemented with complete EDTA-free protease inhibitor cocktail. Cells were lysed by sonication and the supernatant was separated by centrifugation at 18,500x g for 20 min at 4 °C. The soluble fraction was incubated with Ni-NTA resin (Qiagen) for 1 h at 4 °C with mild agitation. Ni-NTA resin was washed before elution with 150 mM imidazole. UBE2A was additionally purified by SEC using a Superdex 75 16/600 column (Cytiva) equilibrated in 50 mM Tris, 150 mM NaCl and 0.5 mM TCEP, pH 7.5.

### Ontology analysis

Differentially enriched GO terms were obtained using online Enrichr ontology analysis tool^98–100^ with GO Biological processes 2023 library or Reactome 2022 library ^101,102^ as a reference. The enriched terms are plotted based on their adjusted p-value and the number of genes within each GO Biological Processes 2023 or Reactome 2022 term. The adjusted p-value is calculated using the Benjamini Hoochberg method for multiple hypotheses testing correction.

## Notes

### Competing Interest Statement

The authors have declared no competing interest.

